# Diurnal cycles drive rhythmic physiology and promote survival in facultative phototrophic bacteria

**DOI:** 10.1101/2023.09.27.559767

**Authors:** Camille Tinguely, Mélanie Paulméry, Céline Terrettaz, Diego Gonzalez

**Affiliations:** Laboratory of Microbiology, Institute of Biology, University of Neuchâtel, Switzerland

## Abstract

Bacteria have evolved many strategies to spare energy when nutrients become scarce. One widespread such strategy is facultative phototrophy, which helps heterotrophs supplement their energy supply using light. Our knowledge on the impact that such behaviors have on bacterial fitness and physiology is, however, still limited. Here, we study how a representative of the genus *Porphyrobacter,* in which aerobic anoxygenic phototrophy is ancestral, responds to different light regimes under nutrient limitation. We show that bacterial survival in stationary phase relies on functional reaction centers and varies depending on the light regime. Under dark-light alternance, our bacterial model presents a diphasic life history dependent on phototrophy: during dark phases, the cells inhibit DNA replication and part of the population lyses and releases nutrients, while subsequent light phases allow for the recovery and renewed growth of the surviving cells. We correlate these cyclic variations with a pervasive pattern of rhythmic transcription which reflects global changes in diurnal metabolic activity. Finally, we demonstrate that, compared to either a phototrophy null mutant or a bacteriochlorophyll *a* overproducer, the wild type strain is better adapted to natural environments, where regular dark-light cycles are interspersed with additional accidental dark episodes. Overall, our results highlight the importance of light-induced biological rhythms in a new model of aerobic anoxygenic phototroph representative of an ecologically important group of environmental bacteria.

## Introduction

In natural environments, free-living bacteria face, between short intervals of fast multiplication, long periods of stasis or very slow growth [1–3]. To survive such periods, they have evolved diverse strategies to optimize resource utilization and spare energy [4–7]. Facultative photoheterotrophy is such a strategy that a subset of environmental bacteria use to supplement their mostly heterotrophic growth. It can be based on two main molecular mechanisms: retinalophototrophy, which uses rhodopsins to pump ions through the membrane, and chlorophototrophy, which is based on photochemical reaction centers containing chlorophylls or bacteriochlorophylls [8]. <u>A</u>erobic <u>An</u>oxygenic <u>P</u>hototrophy, hereafter AAnP, is a common chlorophototrophic pathway, whereby usable energy is produced from light under aerobic conditions in the absence of H_2_O dissociation and O_2_ release [9, 10].

Aerobic anoxygenic phototrophic bacteria include, among others, members of the alpha­, beta­ and gamma­proteobacteria [11]. These represent an important fraction of the prokaryotes present in the upper layers of seas and lakes, but are also widespread in environments as diverse as the phyllosphere, soil crusts, or thermal springs [12–14]. These bacteria encode a set of about 45 genes required to produce bacteriochlorophyll *a,* the main pigment needed for phototrophy, and the components of their light harvesting complexes, including carotenoids [15]. While these bacteria are in general not able to grow photoautotrophically, with the notable exception of *Dinoroseobacter shibae* which can use the ethylmalonyl-CoA pathway [16], they can benefit from light exposure to maintain higher concentrations of intracellular ATP, incorporate limited amounts of CO_2_ via substrate co-assimilation or anaplerotic reactions, and spare organic carbon through downregulation of respiration [17–21]. However, bacteriochlorophyll *a* biosynthesis and AAnP represent a significant energy investment and are known to generate toxic byproducts, especially singlet oxygen and other reactive oxygen species, in the presence of oxygen and light [11, 22, 23]. That is why many aerobic anoxygenic phototrophs tend to regulate bacteriochlorophyll *a* production strictly and limit its synthesis to dark phases [24, 25].

The life history of aerobic anoxygenic phototrophs is likely to be deeply affected by diel cycles, both because these bacteria can actively utilize light when needed and because they tend to associate and cooperate with obligate photoautotrophs [23, 26–29]. While we know the preferences of some species in terms of light intensity and adaptations to non-optimal light conditions [30, 31], our understanding of the rhythmic physiology of these bacteria in response to diurnal cycles and more generally of the adaptive value of AAnP under different light regimes is still limited [32]. Indications can be found scattered in literature pertaining to quite diverse microbiological fields. Field studies in microbial ecology suggest that the transcription of AAnP genes may vary throughout the day, being maximal at night, and that cell division and cell activity are also driven by diurnal cycles in AAnP-competent bacteria [33–35]; in freshwater natural communities grown *ex situ,* respiration rates and organic substrate assimilation can vary in response to periodic infrared light exposure, suggesting that these variations are driven by AAnP [36]. Laboratory based studies of AAnP models show clear differences in global transcription patterns between cells cultured in the dark and in the light, some of these differences being directly connected to light-stress responses [37, 38]. Finally, studies on pure cultures of *Roseobacter* sp. OCH 114 and *Dinoroseobacter shibae* DFL12 under controlled conditions showed that these bacteria benefit from exposure to continuous light or dark-light cycles for their survival, although this survival benefit was never genetically linked to phototrophy [17, 39]. Despite these convergent indications, consistent evidence on a single experimental model, ranging from genetics to fitness measurements, is actually missing. Moreover, it is generally unclear how AAnP affects bacterial fitness under diel cycles and how facultative aerobic anoxygenic phototrophs navigate the trade­off between costs and benefits of phototrophy.

Here, we investigate how a new freshwater model of AAnP-competent bacteria, *Porphyrobacter* sp. ULC335, responds to and benefits from diel dark-light cycles. When grown to stationary phase under dark-light alternance, *Porphyrobacter* sp. ULC335 presents rhythmic patterns of density variation that were not reported in most other species capable of AAnP. We explain these patterns by connecting them to variations in mortality and reproduction rates due to phototrophy, study their transcriptional correlates, and reveal their impact on fitness. In so doing, we contribute to a better understanding of the rhythmic patterns governing the physiology and lifestyle of bacteria prevalent in photic aquatic layers.

## Methods

Detailed methods are available in a “Supplementary Methods” file.

### Standard growth conditions

*Porphyrobacter* sp. ULC335 was grown in BG11 media [40] supplemented with 0.5g/L Tryptone, 0.25g/L Yeast Extract, and vitamins. Standard batch growth took place at 30°C with agitation. Growth in the Photon Systems Instruments MC1000 took place at 28°C. Batch cultures for competition and survival experiments were done in 24-well plates at 28°C with agitation.

### Pigment characterization

Cell pellets were extracted with 7:2 acetone-methanol for 5 minutes; the absorbance of the supernatant was measured in quartz cuvettes (Genesys 10s, Thermo Scientific) or in 96-well plates (SpectraMax i3x, Molecular Devices). For Thin Layer Chromatography, 100µl of extract were spotted onto a Silica 60 F_254_ plates (Merk, art. 5554). The plate was migrated in 8:0.75:0.2:0.8:0.25 petroleum ether:hexane:isopropanol:acetone:methanol.

### Transposon library

pRL27 [41] was conjugated into *Porphyrobacter* sp. ULC335 and several mutants whose color differed from the wild type were isolated. The insertion sites were identified using the method previously published [41]. Four mutants are central to our study: mutant B^+^C^-^ (insertion in *crtI*, coding for the phytoene synthase, required for carotenoid production), mutant B^-^C^+^ (insertion in *bchH*, coding for a subunit of the magnesium chelatase, required for Bchl*a* synthesis), mutant B^-^C^-^ (insertion in *ispA*, coding for the farnesyl diphosphate synthase, required to synthesize a precursor of both Bchl*a* and carotenoids), mutant B^++^C^++^ (insertion in the *ppaA* gene coding for an AAnP regulator).

### Genome sequencing and annotation

The genomic DNA was purified using the Qiagen Genomic-tip 20/g kit and sequenced on PacBio Sequel instruments. The genome was assembled using PacBio *CCS* 4.1.0 for the read correction step and *Flye* 2.6 [42] for the assembly. The functional classification of the genome content was done using the online *blastKOALA* service with the genus *Porphyrobacter* as a reference.

### RNA-sequencing

RNA was extracted from 1 ml culture using TRIzol (Invitrogen) followed by purification on RNeasy mini-kit columns (Qiagen). RNA-sequencing was carried out at Novogene on Illumina NovaSeq systems after ribosomal depletion and in-house library construction. Reads were trimmed and filtered using *fastP* [43], mapped using *bowtie2* [44], and counted using *featureCounts* [45]. Comparison between wildtype and B^-^C^+^ strains was done using the *DESeq2* package [46]. Temporal analysis of individual genes was done using the *ImpulseDE2* package [47]. Each temporally regulated gene was assigned to a cluster using *hclust* (Euclidian algorithm, 15 clusters) on the Z-scores of the *rlog*-transformed count data for all four time points. The *ggVennDiagram* [48] and *ggplot* [49] libraries were used to plot the summary figures.

### Phylogenetic analysis

Fourteen protein sequences characteristic of the AAnP system from complete alphaproteobacterial genomes were concatenated and aligned using *muscle* 3.8.31 [50]; a phylogenetic tree was constructed using *iqtree* 2.1.1 [51] with default parameters (evolution model chosen automatically). In parallel, a set of conserved ribosomal proteins, elongation factors, and gyrases were retrieved and a phylogenetic tree was constructed following the same procedure. Strains were considered to encode the anoxygenic phototrophy gene cluster if they gave positive *blastp* matches with at least 13 out of the 14 proteins.

### Flow cytometry and fluorescence microscopy

Cells stained with LIVE/DEAD BacLight (Invitrogen) were analyzed by flow cytometry (BD Accuri C6 Plus). The zones corresponding to the dead, compromised, and live subpopulations were determined empirically based on the density distribution of live and 2.5% glutaraldehyde-fixed cells. Visualization and quantification was done with *R* [52] using *flowCore* [53]. The same cells were analyzed using fluorescence microscopy, where only dead and live subpopulations could be distinguished.

Cells stained with DyeCycle Orange (Invitrogen) were analyzed by flow cytometry. The coordinates of the highest density in the regions of the 1N and 2N peaks (green fluorescence) were determined empirically. The lowest density point between the 1N and the 2N peaks on the empirical density plot was used as a proxy for the “actively replicating” population.

### Survival and competition experiments

For survival experiments, late exponential phase cultures of wild type, B^-^C^+^, and B^++^C^++^ were diluted and grown under continuous light, continuous dark, or 12h/12h dark-light cycles in triplicates. At the beginning of stationary phase (48h) and after four additional days (144h), CFUs were determined and the survival rate (144h vs 48h) was calculated.

For competition experiments, exponential phase cultures of the two mutant strains were mixed with the wild type strain in equal proportions and grown under continuous light, continuous dark, or 12h/12h dark-light cycles in triplicates. The dual cultures were plated at 48h and 144h like for the survival experiment, and the ratios of the mutant to the wild type were calculated for both timepoints based on CFUs.

In survival and competition experiments, irradiance during light periods was 80 µmol/s/m^2^.

## Results

### Strain ULC335 is a representative of the *Porphyrobacter* genus, in which AAnP is ancestral

In the course of a study on a bacterial community associated with a single cyanobacterial species (ULC335) [54] (SFile 1), we isolated a nonmotile dark-red colony forming bacterial strain which, based on its 16S RNA sequence, was assigned to the *Porphyrobacter-Erythrobacter* group (Erythrobacteraceae genus I-VI) [55]. The strain encoded an allele of the *pufM* gene, coding for a light-harvesting complex subunit, and readily produced, under standard aerobic culture conditions, bacteriochlorophyll *a* (hereafter “Bchl*a*”) (SFig. 1 and 2), suggesting that the strain was capable of AAnP [56, 57].

Bacteria belonging to the *Porphyrobacter-Erythrobacter* group are ubiquitous in aquatic environments and often found associated with Cyanobacteria [58–61]. While many genera among Alphaproteobacteria contain members capable of anoxygenic phototrophy (SFig. 3), the *Porphyrobacter-Erythrobacter* group is one of the few taxonomic units in which the AAnP gene cluster is most likely ancestral and present in the vast majority of sequenced strains (Fig. 1A, SFig. 4). Moreover, very few members of the *Porphyorbacter-Erythrobacter* have lost the gene cluster, which indicates that it consistently provides a benefit. Noteworthy, the dominant mode of transmission of the genes coding for anoxygenic phototrophy within alphaproteobacteria, including within the *Porphyorbacter-Erythrobacter* group, is vertical, suggesting that the gene cluster has extensively co-evolved and co-adapted with the core genome (SFig. 5, SFig. 6). On this basis, we consider the *Porphyrobacter-Erythrobacter* group as an excellent model to study the effects of long-term adaptation to AAnP.

**Figure 1:**
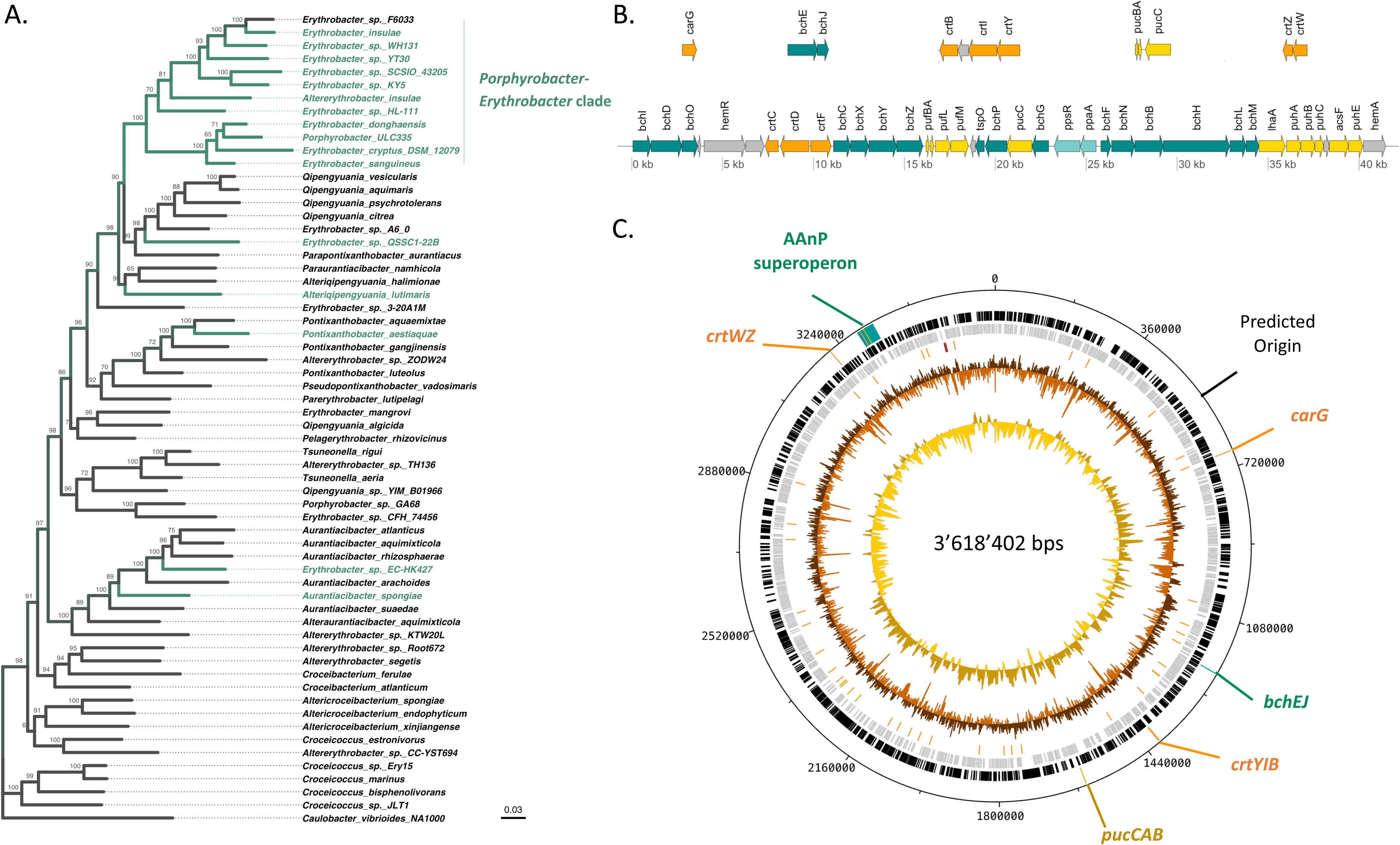
*Porphyrobacter* sp. ULC335 is an aerobic anoxygenic phototroph. A. Simplified phylogenetic tree made from an alignment of concatenated core-genome proteins of Erythrobacteraceae rooted on *Caulobacter crescentus* (Caulobacterales); a tree comprising more strains can be found in SFigure 4. *Porphyrobacter* sp. ULC335 branches within the *Porphyrobacter-Erythrobacter* clade, in which the anoxygenic phototrophy gene cluster is quasi-ubiquitous (branches in green). Scale bar: average number of substitutions per site. Numbers at the nodes are support values (percent of generated trees supporting the branch). B. The aerobic anoxygenic phototrophy (AAnP)-associated genes found in *Porphyrobacter* sp. ULC335. The AAnP-associated genes are scattered in six loci, one major locus more than 40kb in length, and five minor loci, comprising two or three genes. The color codes indicate the predicted functional categories of the products: dark green for proteins involved in bacteriochlorophyll *a* synthesis, yellow for structural proteins of the reaction centers, orange for proteins involved in carotenoid synthesis, light green for regulatory genes, grey for genes associated with the AAnP genes but whose function is not clear. C. Schematic overview of the genome of *Porphyrobacter* sp. ULC335. The concentric circles represent, from inside out, the GC skew (yellow and gold), the GC content (light and dark brown), RNAs (red), genes encoded on the reverse and forward strand (grey and black), AAnP-related loci (same color code as in B), genome coordinates using the coding sequence of *dnaA* as a reference.

To further characterize *Porphyrobacter* sp. ULC335, we sequenced its genome, obtaining a single closed circular sequence of 3.6 Mbps. Besides the typical genetic complement of free-living Alphaproteobacteria (SFig. 7AB), it encoded a complete AAnP gene cluster comparable to the one found in sequenced members of the *Porphyrobacter-Erythrobacter* group (Fig. 1B) [15] and no less than five blue-light responsive proteins including a BLUF domain [62, 63] (SFig. 7C). Some of the genes coding for the carotenoid synthesis pathway are found in the AAnP gene cluster (*crtC, crtD, crtF*), others in secondary loci (*crtZ, crtW; crtB, crtI, crtY; crtG)* (Fig. 1BC). Thin layer chromatography revealed that, besides Bchl*a* and its intermediates, *Porphyrobacter* sp. ULC335 synthesizes a number of red and yellow pigments (SFig. 8AB). *Porphyrobacter* sp. ULC335 does not encode a full carbon fixation pathway and is unlikely to grow autotrophically. However, like *Erythrobacter* sp. NAP1, it produces key enzymes of the 3-hydroxypropionate bicycle pathway for carbon fixation, which would give it the capacity to co-assimilate CO_2_ while using glycolate, glyoxylate, or acetate, among others, as carbon sources (SFig. 9) [19, 64]. Altogether, these data suggest that *Porphyrobacter* sp. ULC335 could make use of light to spare organic substrates under nutrient limitation [20].

### In stationary phase, *Porphyrobacter* sp. ULC335 presents light-induced rhythmic growth patterns dependent on a functional photosystem

When monitored under 12h/12h dark-light cycles, *Porphyrobacter* sp. ULC335 presented, during stationary phase, strong variations in optical density in response to light in a range of light intensities (Fig. 2A, SFig. 10). The bacterial density increased during the light phase and decreased during the dark phase; the rhythms strictly followed the photoperiod and did not persist under constant light or darkness (Fig. 2A, SFig. 11), which suggests that they were not driven by a circadian clock but by external light variations. We hypothesized that such variations in optical density could be a consequence of the phototrophic activity of the organism, although, to our knowledge, such conspicuous rhythms have not been reported so far in bacteria capable of AAnP in the absence of additional phototrophic systems. To test this hypothesis, we generated a Tn5 transposon mutant library of the strain and screened for variations in color which would betray an alteration of Bchl*a* or carotenoid content. We obtained four mutants of interest (Fig. 2B): strain B^+^C^-^ produced Bchl*a* but no carotenoids (insertion in *crtI*, coding for the phytoene synthase), strain B^-^C^+^ produced carotenoids, although less than the wild type, but no Bchl*a* (insertion in *bchH*, coding for a subunit of the magnesium chelatase required for Bchl*a* synthesis), strain B^-^C^-^ produced neither Bchl*a* nor carotenoids (insertion in *ispA*, coding for the farnesyl diphosphate synthase, required to synthesize a precursor of both Bchl*a* and carotenoids), strain B^++^C^++^ overproduced both Bchl*a* and carotenoids, especially in rich medium and under continuous light (insertion in the *ppaA* gene coding for an AAnP regulator) (Fig. 2C; SFig. 12). Of the four mutants, only the B^++^C^++^ strain presented rhythmic growth similar to the wild type strain (Fig. 2D). By contrast, strains B^-^C^-^ and B^-^C^+^ did not present any periodic optical density variation (Fig. 2E). The strain B^+^C^-^ was very sensitive to light: as soon as light was switched on the density started decreasing and continued during the subsequent cycles (Fig. 2E); upon light exposure, the viability of cells, as quantified by Life/Dead staining and CFUs, decreased by at least two orders of magnitude (SFig. 13). This suggests that, in the absence of carotenoids, Bchl*a* production might be toxic to the cells, especially in the presence of light. Overall, these data indicate that the rhythmic density variations in response to light depend on a functional reaction center including both Bchl*a* and carotenoids.

**Figure 2:**
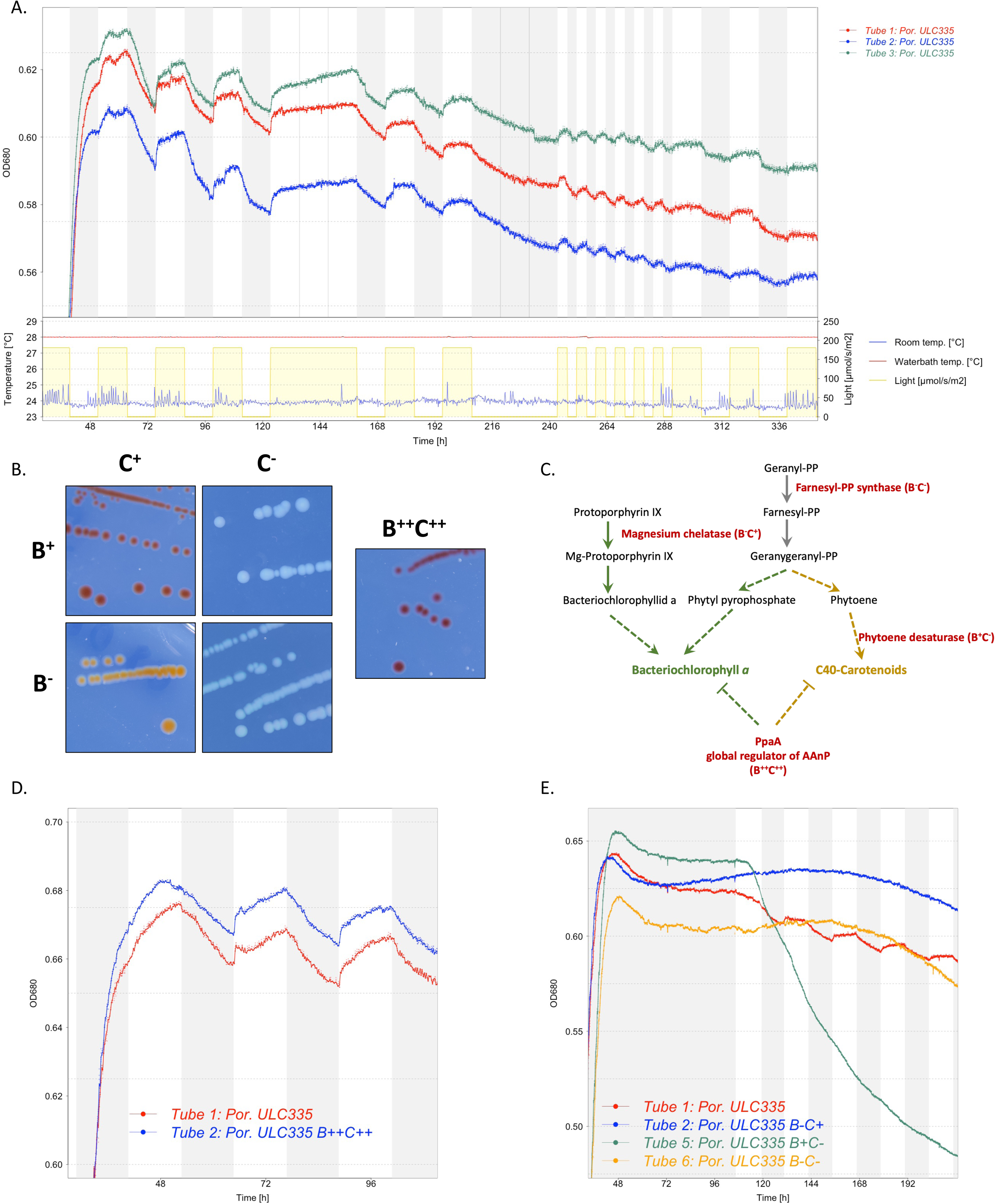
*Porphyrobacter* sp. ULC335 presents rhythmic growth patterns in stationary phase that are dependent on functional anoxygenic phototrophy. A. Rhythmic variations in optical density (measured at 680nm) of three replicate cultures of *Porphyrobacter* sp. ULC335 under dark-light alternance (light at 180µmol/s/cm^2^). Light intensity, culture temperature (water bath), and room temperature are shown as evidence that density changes are from biological and not technical origin. Cultures were maintained in 80 ml tubes in a device adapted for the culture of cyanobacteria or algae. The rhythms do not present persistence under continuous light (120h-156h) or continuous darkness (204h-240h), and follow light variation seamlessly when the period is shorter than 24h (240h-288h). B. Colony phenotype of *Porphyrobacter* sp. ULC335 (B^+^C^+^) and four transposon mutants grown on BG11-P-agar in the dark. The color of colonies are in agreement with the absorbance profiles of liquid cultures (SFigure 10). C. Simplified pathway of carotenoid and bacteriochlorophyll *a* biosynthesis. In red, the genes in which the transposon is inserted for each of the mutant strains. Ticked lines represent multiple synthesis steps (enzymatic reactions) or multiple convergent effects (regulators). D. Rhythmic variations in optical density (measured at 680nm) of the wild type compared with the bacteriochlorophyll *a* and carotenoid overproducer strain. No significant difference in pattern is apparent between the two strains. The culture conditions are similar to panel A. E. Variations in optical density (OD, measured at 680nm) of the wild type compared with the three bacteriochlorophyll *a* and/or carotenoid mutant strains. B^-^C^+^ and B^-^C^-^ did not present any rhythmic growth pattern; B^+^C^-^ presented slight variations in slope between dark and light phases, the speed of the OD decline increasing during light phases. Because B^+^C^-^ was strongly inhibited by light, the cultures were maintained under continuous darkness until stationary phase, when a 12h/12h dark-light alternance illumination regime was started.

### The optical density variations in response to light correlate with changes in cell viability and DNA replication

To further investigate the origin of these rhythms, we sought to determine what happens to *Porphyrobacter* sp. ULC335 cells when they are exposed to darkness. To do so, we sampled six parallel cultures of *Porphyrobacter* sp. ULC335 grown to stationary phase and maintained under continuous light for two days; three of the cultures were switched to the dark for 12 hours before sampling (Time Point 1 [TP1]) and switched back to light immediately after sampling (dark treatment); a second sample of all six tubes was taken eight hours after the switch back to light (TP2) (continuous light treatment) (Fig. 3A, SFig. 14). Using spectrophotometry and flow cytometry, we did not find any difference in pigment content, cell morphology or aggregation between the treatments (SFig. 15). However, using Live/Dead staining, we observed a clear difference in the proportion of living and compromised (permeable) cells between the two treatments: while 70% cells cultured under continuous light were viable and non-compromised, based on fluorescence microscopy, this proportion dropped to 40% after 12 hours in the darkness; interestingly, it was enough for the latter cultures to be exposed back to light for 8 hours to restore the proportion of viable cells to original levels (Fig. 3B). Colony forming units confirmed this trend (SFig. 16). Flow cytometry revealed that three subpopulations coexist in the cultures, independently of the treatment: viable noncompromised cells, compromised (permeable) cells, and dead cells (Fig. 3C; controls: SFig. 17). Like the “dead” subpopulation, the “compromised” subpopulation was significantly larger in the cultures exposed to 12 hours of darkness than in the cultures exposed to continuous light (SFig. 18). Moreover, by analyzing the DNA content of the cells, we found that the frequency of cells undergoing DNA replication was significantly lower in the cultures exposed to darkness than in the cultures kept under continuous light; upon reexposure to light, the two populations were indistinguishable again (Fig. 3D; SFig. 19). Taken together, these results suggest that the life history of *Porphyrobacter* sp. ULC335 follows alternate phases in response to illumination conditions: during dark-phases, cells slow down replication and a significant proportion of them dies; during light-phases, part of the surviving cells replicate and divide, and this, possibly along with the partial recovery of some compromised cells, leads to the restoration of previous viability levels in the population.

**Figure 3:**
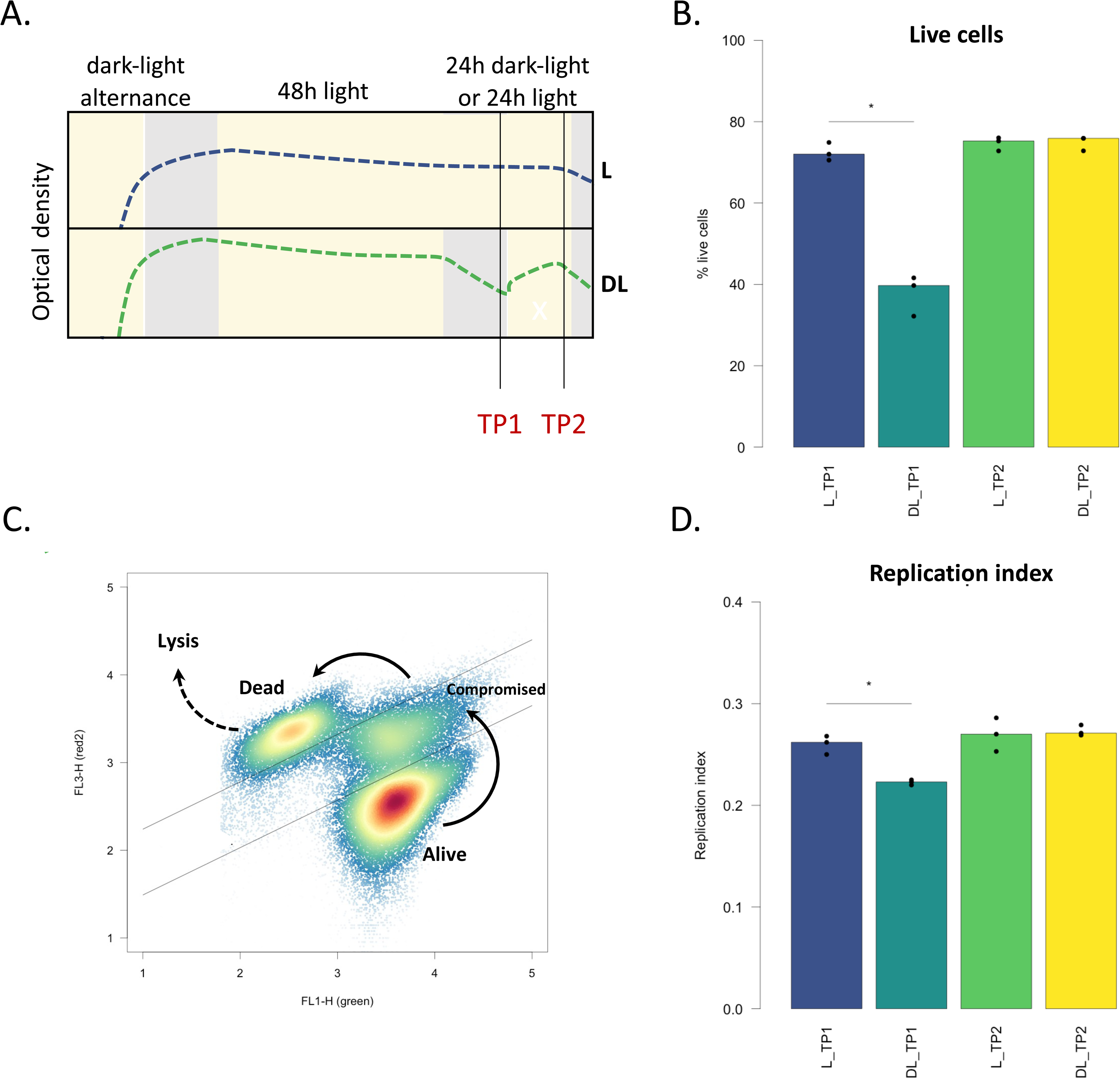
Variations in optical density correlate with differences in cell death and replication rates. A. Experimental design. Six *Porphyrobacter* sp. ULC335 cultures were maintained in parallel under dark-light alternance up to stationary phase to allow for carotenoid and bacteriochlorophyll *a* accumulation; then, the cultures were exposed for 48 hours to continuous light, before three of the six cultures were switched to the dark for 12 hours and back again to light for another 12 hours. Two series of samples were taken, one at the end of the dark period and one 8 hours after return to light, and analyzed using fluorescence microscopy and flow cytometry. B. Living cells as determined using fluorescence microscopy on Live/Dead stained cells. The proportion of living cells dropped after the dark phase and returned to baseline after exposure to light. At least 500 cells were counted per replicate. C. Dead, compromised, and alive subpopulations as observed using flow cytometry after Live/Dead staining. The arrows indicate the probable path from alive to compromised to dead cells, as membrane permeability increases; lysed cells have released their cellular content, including DNA, and cannot be observed using flow cytometry. D. The proportion of replicating cells, as determined by flow cytometry using DyeCycle Orange stain, decreased during the dark phase and returned to baseline after reexposure to light. A total of 40’000 cells was analyzed. The calculation of the replication index is explained in the methods; see Sfigure 17 for a more detailed representation of flow cytometry data. Amount of light during light periods: 150 µmol/s/m^2^. Asterisks indicate statistically significant differences (bilateral Student’s t.test, p-value<0.05).

### In *Porphyrobacter* sp. ULC335, direct responses to light and responses to electron flow underlie rhythmic physiology

To understand the physiological correlates of this behavior, we sequenced the full transcriptome of stationary phase cultures under 12h/12h dark­light cycles. We were particularly interested in transcriptional responses upon light phase transition, so we sampled one hour before (Late Night, “LN”) and after (Early Day, “ED”) the switch to light, one hour before (Late Day, “LD”) and after (Early Night, “EN”) the switch to darkness. In addition, we treated in parallel the wild type strain and the arrhythmic phototrophy­mutant B^-^C^+^ in order to be able to discriminate between direct responses to light (independent of Bchl*a*) and responses due to the electron flow generated by phototroph.

Disrupting photosynthesis had massive consequences at the transcriptional level. Among protein-coding genes, 1533 out of 3418 were differentially regulated more than two-fold at least at one of the four time points in the phototrophy mutant, about half being repressed and half being activated (Fig. 4A; SFile 2). The time of day had a strong influence on the divergence of the transcriptional profile between the two strains. The number of differentially regulated genes between the two strains differed the most at the EN time point and differed the least at the LN timepoint, suggesting that, in the prolonged absence of light exposure, the physiologies of the two strains tended to converge. Globally, the mutant and the wild type seem to time transcription differently. Genes more transcribed in the mutant compared to the wild type were most abundant during the day (and EN), while genes less transcribed in the mutant compared to the wild type were most abundant during the night. Among genes repressed in the mutant, functions linked to photosynthesis, including porphyrin metabolism, and ribosome biogenesis were enriched, especially during the day; genes activated in the mutant included categories linked to DNA repair, translation, and energy metabolism (Fig. 4B; SFile 3). This indicates that the transcriptional investment into energy metabolism, especially photosynthesis, and translation machinery is modified, both in terms of timing and intensity, in the phototrophy null mutant compared to the wild type.

**Figure 4:**
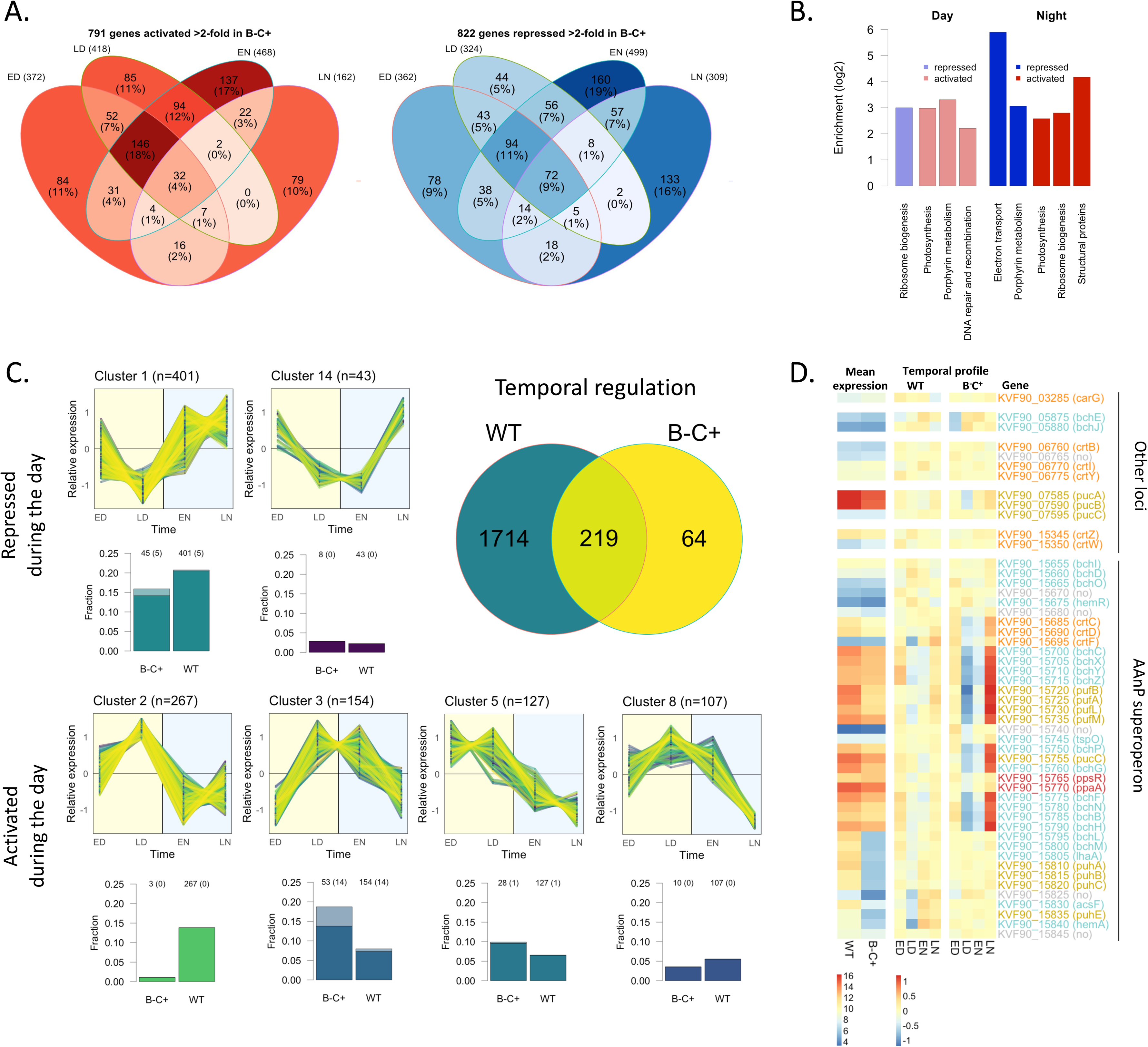
Diel dark-light cycles drive extensive transcriptional rhythms in *Porphyrobacter* sp. ULC335. A. Summary of the experiment: number of genes activated (right) or repressed (left) in the Bchl*a* null mutant compared to the wild type at different timepoints (ED: early day; LD: late day; EN: early night; LN: late night). B. KEGG categories enriched among differentially regulated genes in the Bchl*a* null mutant compared to the wild type; ED and LD on one side and EN and LN on the other side are considered together in this analysis. C. The figure summarizes information about the temporal regulation of genes in the wild type strain and its Bchl*a* null mutant. The Venn diagram indicates the number of temporally regulated genes in the two strains. The major clusters of genes showing a consistent light-response pattern in either of the two strains are presented based on their temporal profile (upper plot; the datapoints represent the relative transcription rates [the Z-scores of the rlog-transformed count data]) and the fraction they represent among temporally regulated genes in both strains (lower plot; light colors represent genes falling in the same cluster in the wild type and the Bchl*a* null mutant). The same data split by strain is presented in SFig. 22). D. Heatmap of gene expression for the AAnP-associated genes. The first two columns show the relative expression of each gene in the two strains; the next eight columns represent the variation over time of the two strains. The gene number and name are shown in cyan for Bchl*a* biosynthesis genes, gold for reaction center genes, orange for carotenoid biosynthesis genes, and red for regulators; grey is used for genes whose function is dubious.

We then identified genes differentially regulated over time (Impulse DE algorithm, p-value < 0.01) in either the wild type or the B^-^C^+^ strain, clustered them by temporal profile and compared the different sets of genes between the two strains (SFile 4; SFig. 20). Again, photosynthesis disruption had a huge impact: only 283 genes (8% of the genome) varied over time in the B^-^C^+^ strain, compared to 1933 genes (56% of the genome) in the wild type strain. This indicates that functional AAnP not only provides energy to the cells, but also drives a pervasive rhythmic physiology that is entirely absent in the phototrophy mutant. More than half of the genes temporally regulated in the wild type strain show opposite expression trends during daytime and nighttime. Four gene clusters (655 genes in total) are consistently activated during the day and repressed during the night (Fig. 4C, lower part), while two clusters (444 genes in total) are repressed during the day and activated during the night (Fig. 4C, upper part). Overall, this suggests that the transcriptional profile, and consequently the physiology, of *Porphyrobacter* sp. ULCC335 presents a marked contrast between day and night involving opposed regulation of more than 1000 genes (31% of the genes). Genes repressed during the day and activated during the night are enriched for the AAnP pathway, ribosome biosynthesis, and oxidative phosphorylation, including the electron transfer chain; genes activated during the day and repressed during the night include transporters, energy metabolism and especially genes involved in the 3-hydroxypropionate bicycle and possibly implicated in CO_2_ co-assimilation (SFile 5). This suggests that the night is dedicated to replenish Bchl*a* and the ribosomal content of the cells, while nutrient assimilation might be privileged during the daytime.

Like in other AAnP-competent bacteria, genes involved in Bchl*a* biosynthesis and components of the light harvesting complexes tended to be repressed during the day and activated at night in both the wild type and the Bchl*a* null mutant strain (Fig. 4D; SFig. 21). However, the amplitude of expression changes across the four timepoints was attenuated in the wild type strain compared to the Bchl*a* null mutant. In B^-^C^+^, more than half of the expressed genes presented a four-fold difference in expression between late night (maximal expression) and late day (minimal expression), while a variety of attenuated and shifted profiles was represented in the wild type. These differences in the expression of AAnP-related genes suggest that genes which mainly respond to light cues in the B^-^C^+^ strain are tuned, in the wild type, via mechanisms sensitive to energy flow which are only active when AAnP is functional.

Overall, our transcriptome analysis revealed that diel light cycles have a major impact on the transcriptional program and physiology. The activity of the cells differs markedly between the day time and the night time, and there is a clear physiological shift between early and late timepoints in either day or night. Consistent with our observations on optical density, inactivation of AAnP hampers rhythmic transcription and leads to a transcriptional profile that is most similar to the one presented by the AAnP-competent strain at the end of the night, when the benefits of phototrophy wear off.

### Phototrophy provides a survival benefit in stationary phase that correlates with Bchl*a* production

*Porphyrobacter* sp. ULC335 cultures present oscillations in density in stationary phase that correlate with changes in mortality and reproduction rates. However, it is unclear how these variations affect the overall survival of the bacteria, especially in view of the absence of any density variation in Bchl*a* null strains (B^-^C^+^ and B^-^C^-^). We took advantage of the set of mutants we had generated (Fig. 2B-C) to ask whether AAnP actually provides some fitness advantage to *Porphyrobacter* sp. ULC335 depending on the illumination regime (DL: 12h/12h Dark-Light alternance, D: constant Dark, L: constant Light).

While AAnP did not provide a significant advantage in exponential up to early stationary phase (SFig. 22 and 23), survival after 96h spent in stationary phase was strongly affected by the genotype and the illumination regime. This is consistent with the accumulation pattern of Bchl*a* and carotenoids, whose content increases a lot between 24h and 48h of growth, especially in the D and DL treatments, in the two strains capable of AAnP (Fig. 5A). When AAnP was impossible, either in the absence of light or in the absence of Bchl*a* (B^-^C^+^ mutant), the CFUs decreased by about two orders of magnitude leading to survival rates around or below 5% (Fig. 5B). The capacity to produce energy through AAnP helped the wild type and B^++^C^++^ strains survive better under the DL regime, reaching 35% and 60% survival respectively. Under continuous light, again, the wild type and B^++^C^++^ strains survived better than the Bchl*a* null mutant, but mortality was nonetheless much higher than under the DL regime, presumably because Bchl*a* accumulation was impaired (Fig. 5A). These results demonstrate that AAnP provides a survival advantage to *Porphyrobacter* sp. ULC335 that depends on both growth phase and illumination regime, and partially correlates with Bchl*a* accumulation. In our strain, AAnP is therefore not primarily used to supplement active heterotrophic growth, but to enhance survival during periods of starvation, especially under DL illumination.

**Figure 5:**
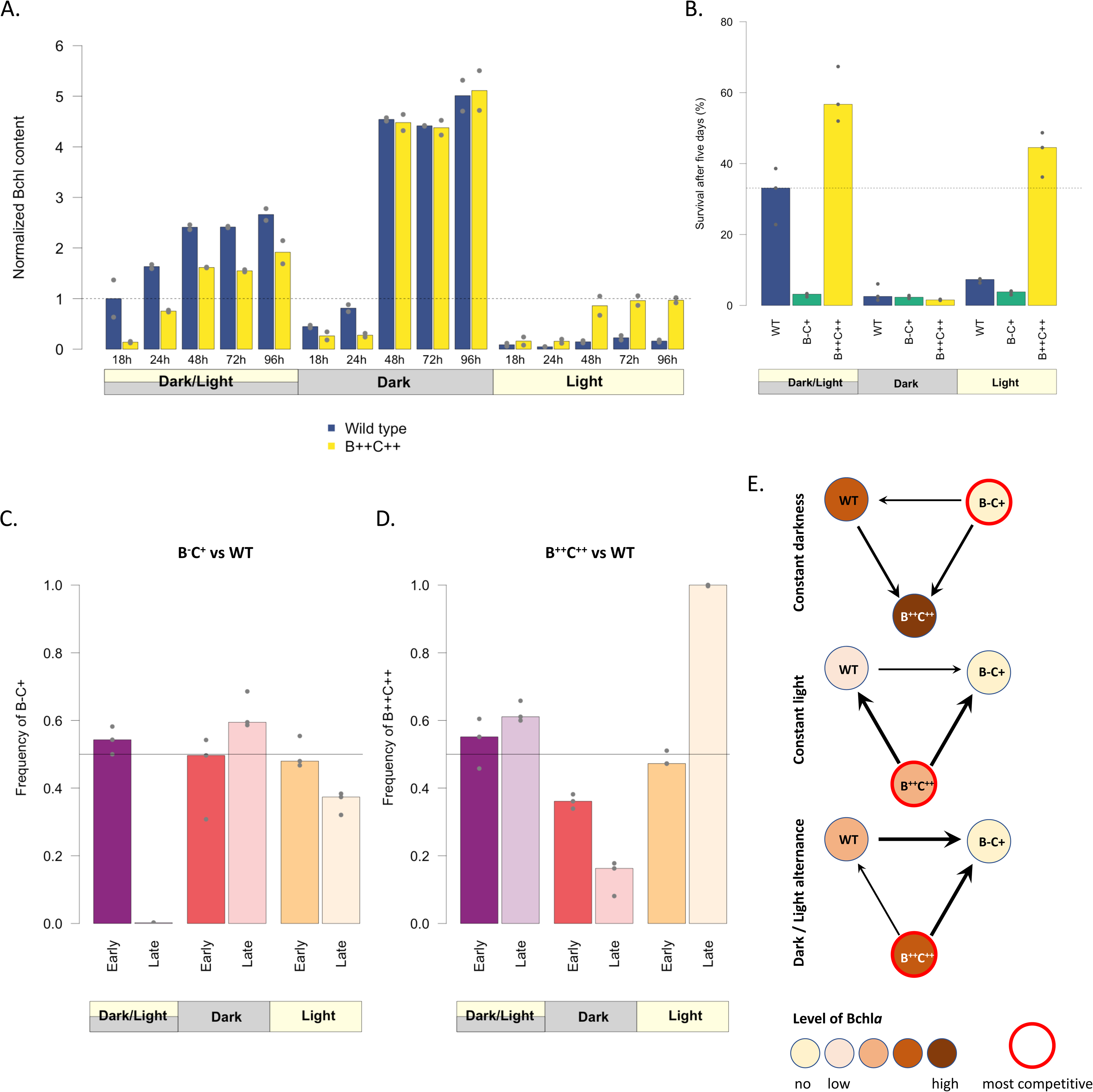
Phototrophy provides a major benefit to *Porphyrobacter* sp. ULC335 in stationary phase. A. Bchl*a* content at different timepoints during batch culture. The barplots represent the absorbance at 770nm (Bchl*a*) corrected by the optical density of the culture (average of two replicates), normalized so that the average of the first timepoint of the wild type strain equals 1. The wild type strain is shown in blue, the Bchl*a* overproducer (B^++^C^++^) in yellow. The experiment was done under 12h/12h dark-light cycles, under continuous darkness, or under continuous light. B. Percent survival, based on colony forming units, of the wild type, the Bchl*a* null mutant (B^-^C^+^) and the Bchl*a* overproducer (B^++^C^++^) after five days in stationary phase. The experiment was done under 12h/12h dark-light cycles, under continuous darkness, or under continuous light. The baseline is taken in early stationary phase (48 hours of growth). C. and D. Outcome of competitions between the wild type and each of the two mutants after five days in stationary phase. The two competing strains were mixed in equal proportions after they had reached stationary phase (48h of growth). E. Figure summarizing the competitivity of each strain under different illumination conditions. The inner color of the circles vary with relative Bchl*a*/carotenoid content; the most competitive strains are outlined in red; the relative competitivity is represented by arrows of three different thicknesses.

To complement these conclusions, we ran competition experiments in which the wild type strain and a competitor strain (either B^-^C^+^ or B^++^C^++^) were grown together in batch under a DL, D, or L regime. We sampled the cultures at 48h (start of stationary phase) and at 144h (late stationary phase), and compared the ratios between the strains. The competitivity of the strains varied depending on the light regime in predictable ways (Fig. 5CD). In the absence of light (D), the Bchl*a* null mutant outcompeted both Bchl*a* producers, and the wild type strain outcompeted the B^++^C^++^ strain. In the presence of light, under both the DL and L regimes, the B^++^C^++^ strain outcompeted the wild type strain which, in turn, outcompeted the Bchl*a* null mutant. Altogether, these results indicate that, in *Porphyrobacter* sp. ULC335, AAnP has a significant cost, including in stationary phase, but that the fitness advantages it provides as soon as light is present—even when light is continuous and prevents the efficient accumulation of Bchl*a*—warrants its maintenance in natural populations.

It stands out that, in our experimental conditions, the wild type strain is less fit than the Bchl*a* and carotenoid overproducer B^++^C^++^ whenever light is present. This suggests that, in the natural environment of *Porphyrobacter* sp. ULC335, the photoperiod is on average lower than 12h or that the regular diel dark-light alternance can often be interrupted by periods of continuous accidental darkness (Fig. 5E).

## Discussion

In this work, we have isolated and described a new *Porphyrobacter* strain, which we found associated with a freshwater cyanobacterium (*Snowella* sp.). In stationary phase, this strain presented rhythmic variations in optical density in response to dark-light cycles, that we correlated with phases of high mortality and low reproduction in the darkness and phases of lower mortality and higher reproduction in the light. This behavior is dependent on Bchl*a* production and a functional reaction center, including carotenoids contributing to photoprotection and light harvesting. Underlying the cycles, we found pervasive rhythmic transcription patterns arising from direct responses to light, possibly driven by any of the five BLUF-domain containing proteins encoded in the genome, and indirect responses to increased electron flow and oxidative stress caused by phototrophic activity. Production of Bchl*a* was found to be costly under continuous darkness and inefficient under continuous light; it provides maximal survival benefits in stationary phase under dark-light cycles, indicating that the strain is adapted to diurnal light cycles. So overall, in this study, we described a distinct bacterial rhythmic behavior, explained it in terms of fluctuations in death and reproduction rates at the cellular level, partially elucidated the transcriptional patterns that underlie it, and linked it to a survival benefit, providing a rationale for its evolution.

Based on average nucleotide identity with known genomes, *Porphyrobacter* sp. ULC335 is likely a new species in the *Porphyrobacter-Erythrobacter* clade (SFig. 24). While *Porphyrobacter* sp. ULC335 probably obtains most of its energy, reductive power, and carbon from organic substrates, it can use phototrophy to supplement heterotrophic growth and to ensure survival when organic substrates become limiting like other AAnP-competent bacteria [11]. Because of the phylogenetic prevalence of AAnP in the clade, these organisms are expected to have evolved ways to tune their physiology to nutrient and light fluctuations, as evidenced by the high numbers of blue-light sensing proteins encoded in the genome of *Porphyrobacter* sp. ULC335 together with homologs for PpsR, AppA, and FnrL, known to integrate phototrophy with metabolism[65, 66]. Because *Porphyrobacter* strains are frequently associated with filamentous or colonial Cyanobacteria [67–69] their physiology could also indirectly follow the light and circadian clock-driven life cycle of their host like it has been suggested for other Cyanobacteria-associated heterotrophs [70]. This makes members of the *Porphyrobacter* genus excellent candidates for the study of bacterial rhythmic behaviors and interactions.

The conspicuous light-driven oscillations in optical density we detected in batch cultures of *Porphyrobacter* sp. ULC335 have never been reported or studied so far in AAnP-competent bacteria, with the exception of *Sphingomonas glacialis* AAP5, recently isolated from an alpine lake, which combines retinalophototrophy and chlorophotoptrophy [71]. In our case, the oscillations involve an increase in density during light phases and a decrease in density during dark phases. Interestingly, the net difference in optical density per 24-hour period is almost null (SFig. 10) suggesting that the biomass lost during dark phases is regained during light phases. The full oscillatory pattern is strictly dependent on both Bchl*a* and carotenoids; however, when Bchl*a* is produced in the absence of carotenoids, light phases trigger a sharp decrease in optical density that hardly slows down during dark phases (Fig. 2E, SFig 13). This suggests that Bchl*a,* or some derivative of it, might become toxic, for instance by generating reactive oxygen species, in the presence of light and cause cell lysis, including in a delayed way during the dark phase [22, 72, 73]. We showed that the decrease in optical density observed during dark phases correlates with an increase in the frequency of membrane-compromised and non-viable cells and a decrease in the frequency of replicating cells in the bacterial population. Based on the decrease in optical density, which corresponds to about 2 to 3% of the total biomass, it is likely that only a fraction of the 20% to 40% of compromised or dead cells (see Fig. 3B) actually lyse during the night. The decrease in compromised or dead cell proportion during the light phases is therefore most likely due to membrane repair and, to some extent, to continued lysis of affected cells, compensated by cell multiplication. Overall, these observations suggest that AAnP functions as a double edge sword: when exposed to light, AAnP photosystems sustain energy production and recycling of organic compounds; during dark phases, however, when energy becomes scarce, a delayed phototoxicity linked to Bchl*a* exposure to light leads to compromised membranes and cell lysis; during the following light phase, the population recovers both in terms of membrane integrity and in terms of viability. It is however unclear whether this recovery is mostly due to compromised cells being able to repair their membrane or to the renewed growth of the surviving population based on nutrients released by lysing cells.

Physiological rhythms in bacteria and especially in heterotrophs remain understudied. Our RNA-sequencing data set reveals that, when AAnP is active, the alternance of dark and light phases affects the transcription of more than 50% of the genes. The transcripts responding to light in the absence of Bchl*a* are only a minority of at most 10%. This indicates that physiological rhythms are pervasive in AAnP-competent bacteria and most likely generated through complex interactions between direct light responses (via light sensors), and indirect responses to light stress, redox and energy state, metabolism, cell-cycle, etc. About half of the genes are consistently activated or repressed by light. This suggests that there is a degree of temporal separation of cellular activities between day and night in these bacteria. Night-activated genes include many genes connected to translation, especially components of the ribosome and translation complexes, as well as genes involved in phototrophy like in other AAnP-competent bacteria [37, 38]. This suggests that dark phases are dedicated to the replenishment of components essential for light harvesting and cellular functioning; they tend to limit, however, active DNA replication as observed through flow cytometry. By contrast, light phases promote the transcription of genes involved in energy metabolism and protection against oxidative light stress, but also key genes in DNA replication (*dnaA* which falls in cluster 2, among genes maximally expressed during the day) and cell division (*ftsZ*, which falls in cluster 4, among proteins maximally expressed by the end of the day); these genes and functions would stimulate DNA replication and cell division, supporting our flow cytometry data (Fig. 3D).

Despite a few precursory studies, the evolutionary benefit of AAnP has not benefited from much attention. Previous research has found that exposure to light increases survival of *Roseobacter* and *Dinoroseobacter* strains in stationary phase, but without formal demonstration that this benefit can be traced back to AAnP [17, 39]. Here we provide, based on manipulation of genotypes and growth environments, an in-depth assessment of costs and benefits of AAnP in terms of its effect on survival and fitness in stationary phase. We find that Bchl*a* production has a significant cost under constant darkness, where Bchl*a* producers get outcompeted by nonproducers. By contrast, under continuous light or alternance, Bchl*a* producers outcompete nonproducers. Under continuous light and, to a lesser extent, under light alternance, strains producing higher levels of Bchl*a* and/or carotenoids get a strong selective advantage. This suggests that the regulation of AAnP in the wild type *Porphyrobacter* sp. ULC335 is optimized for dark-light cycles with a photoperiod lower than 12h or for environments where cells can be accidentally cut off from light by opaque bodies or move out of the photic zone.

## Supporting information

SFile_5

SFile_4

SFile_3

SFile_2

SFile_1

## Acknowledgments

We thank Annick Wilmotte for providing the ULC335 community. We express our gratitude to Pilar Junier, Martina Valentini, and Despoina A. Mavridou for their constructive comments on earlier versions of this manuscript. This work was mainly funded by a Swiss National Foundation Ambizione grant to D.G. (PZ00P3_180142). The contribution of C.Ti. was supported by a grant from the Fondation Mercier pour la Science to D.G.

## Contributions (CRediT)

<u>C.Ti.</u>: Investigation (supporting), Visualization (supporting), Formal analysis (supporting), Review and editing (equal).

<u>M.P.</u>: Investigation (supporting), Review and editing (equal).

<u>C.Te.</u>: Investigation (supporting), Review and editing (equal).

<u>D.G.</u>: Investigation (lead), Conceptualization (lead), Visualization (lead), Data curation (lead), Formal analysis (lead), Project administration (lead), Software (lead), Writing (lead), Review and editing (equal).

## Supplementary Figures

**SFigure 1:**
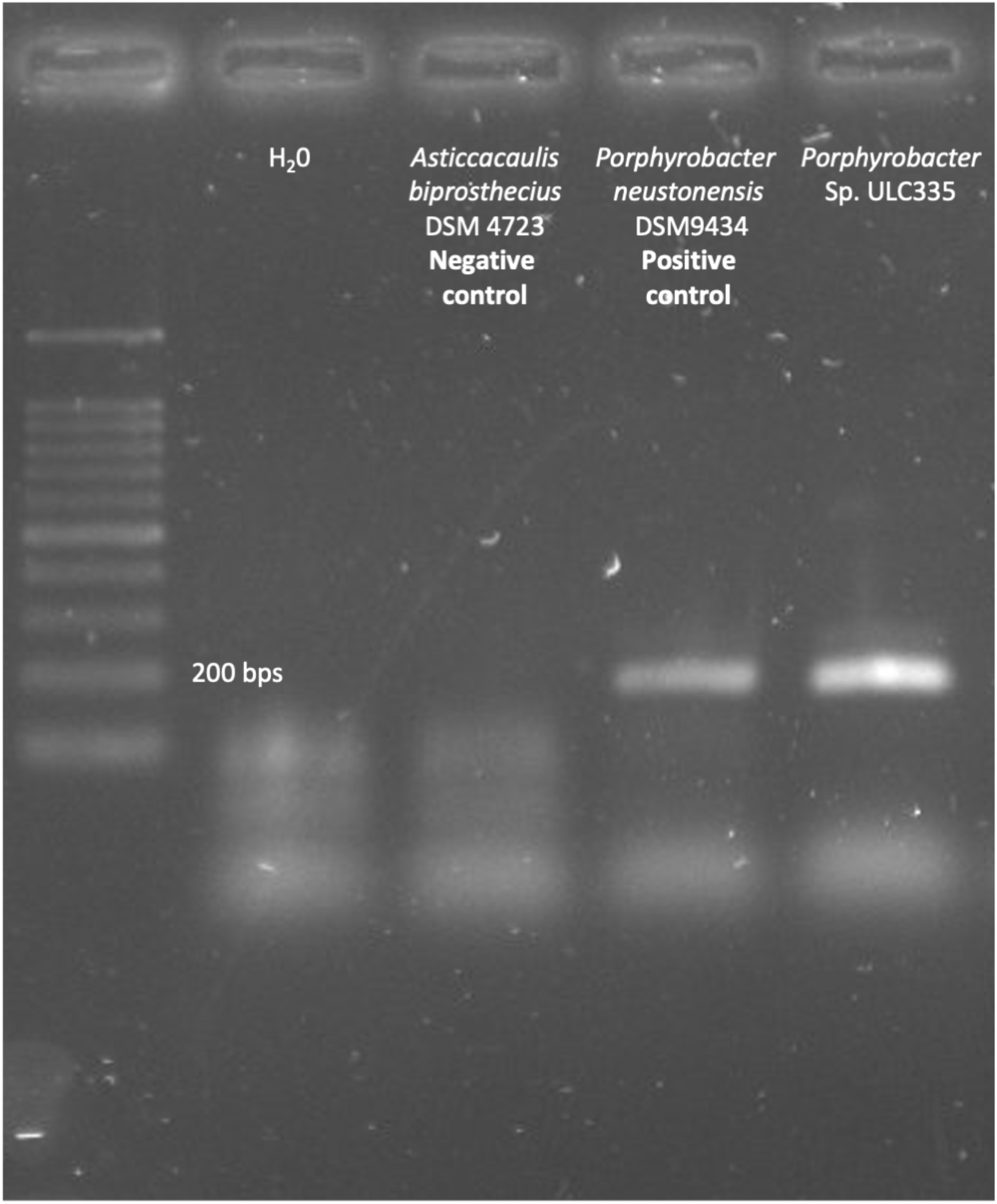
The genome of *Porphyrobacter* sp. ULC335 encodes the *pufM* gene. The figure shows the results of a PCR with primers pufM-F and pufM-R detecting the *pufM* gene (coding for a structural component of the reaction center). Like the type species *Porphyrobacter neustonensis, Porphyrobacter* sp. ULC335 gives a positive signal.

**SFigure 2:**
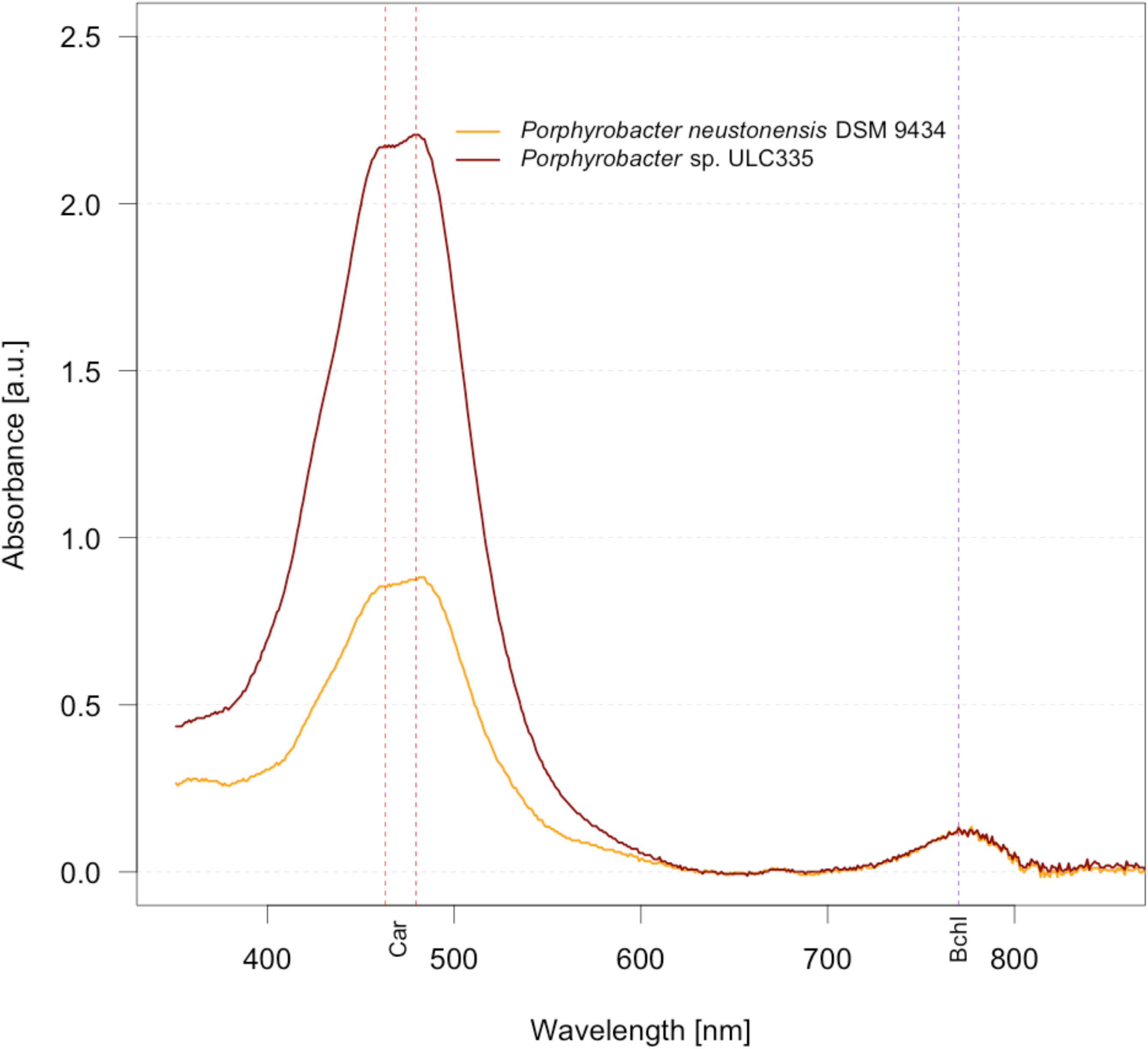
*Porphyrobacter* sp. ULC335 produces pigments whose absorbance is consistent with carotenoids and bacteriochlorophyll. ***a.*** The figure shows the absorbance profile of 7:2 acetone-methanol extracts of 48-hour cultures of *Porphyrobacter neustonensis* and *Porphyrobacter* sp. ULC335. In both species, peaks are found around 460 and 480 nm (carotenoids), and around 770 nm (bacteriochlorophyll *a*). *Porphyrobacter* sp. ULC335 is noticeably more pigmented than *P. neustonensis.* Car: maxima for carotenoids; Bchl: maximum for bacteriochlorophyll *a*.

**SFigure 3:**
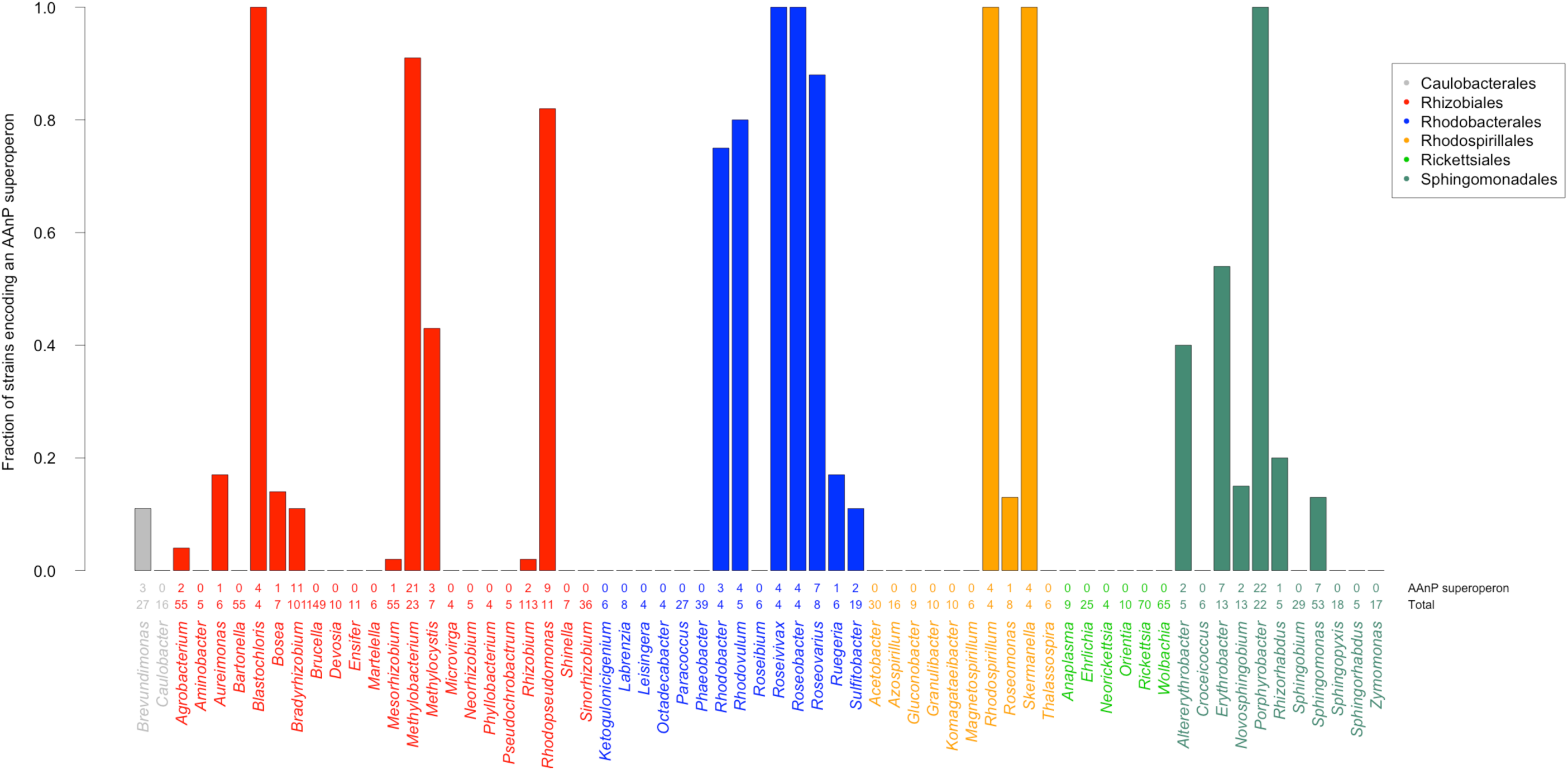
Fraction of genomes per genus within alphaproteobacteria encoding the anoxygenic phototrophy superoperon. Genomes encoding at least 13 out of 14 signature proteins of the anoxygenic phototrophy superoperon were considered positive. Only genera including at least four full genomes were considered.

**SFigure 4:**
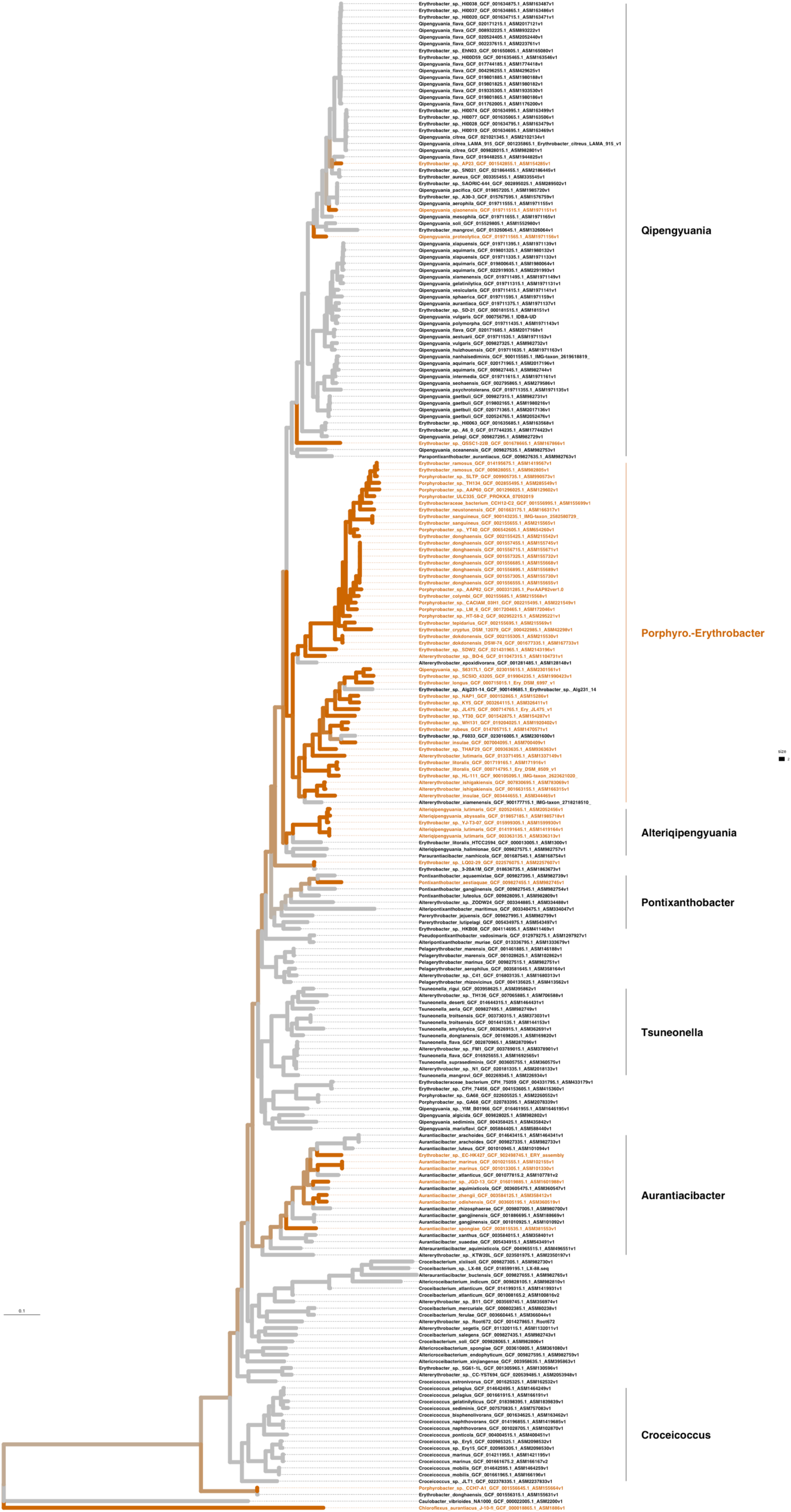
Presence of the anoxygenic phototrophy superoperon within Erythrobacteraceae. The phylogenetic tree was constructed from a set of conserved core proteins from all genomes of Erythrobacterales listed on the NCBI repository. The anoxygenic phototrophy operon was considered present in the genome of at least 13 out of 14 signature proteins were found in it through a *blastp* search. The superoperon is clearly overrepresented in the *Porphyrobacter-Erythrobacter* clade. Scale bar: average number of substitutions per site.

**SFigure 5:**
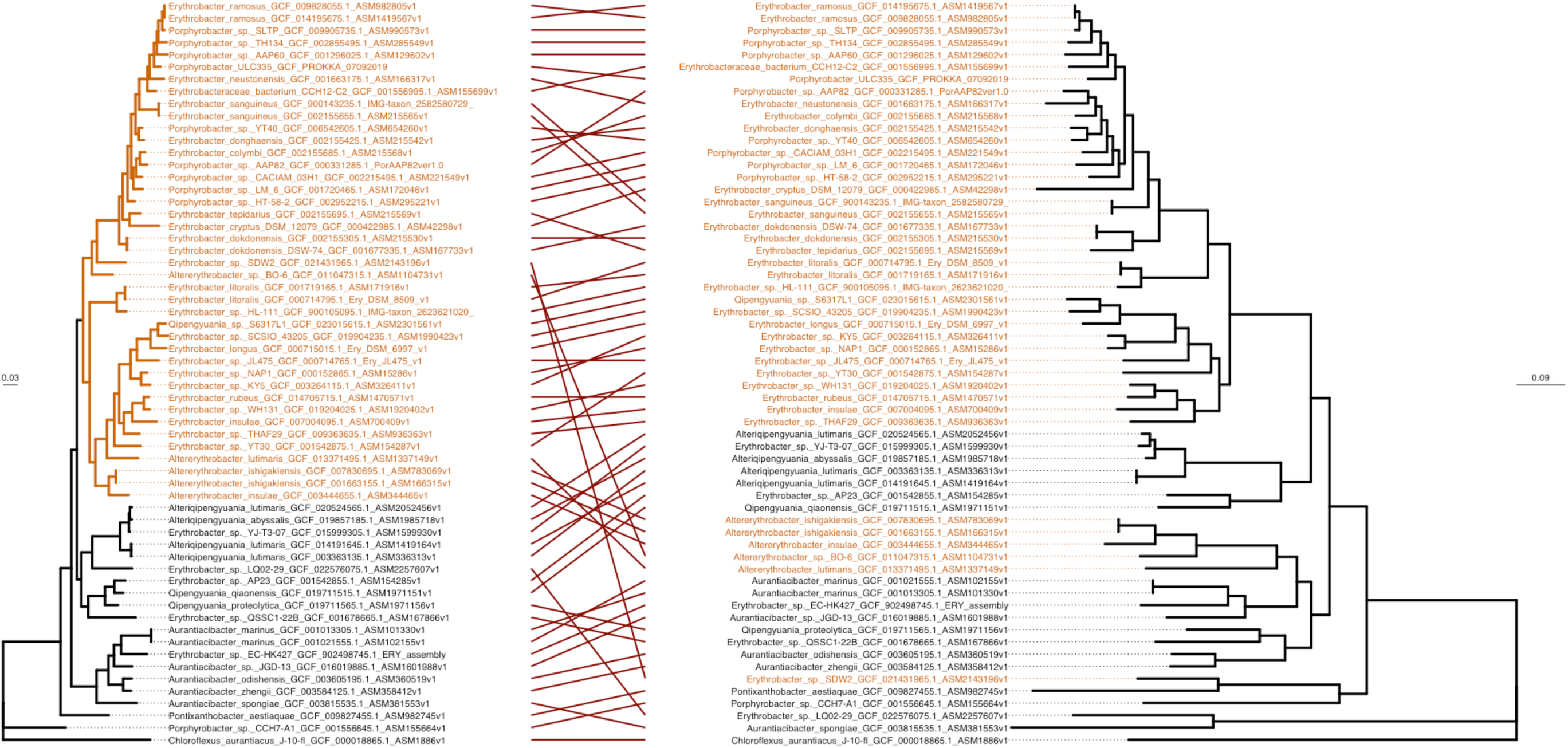
The anoxygenic phototrophy superoperon is primarily transferred vertically in Erythrobacteraceae. In general, the branching order is congruent between a tree made from concatenated conserved core-genome proteins (left) and a tree of concatenated proteins belonging to the anoxygenic phototrophy operon (right). Exceptions are most likely due to horizontal gene transfer. Scale bar: average number of substitutions per site.

**SFigure 6:**
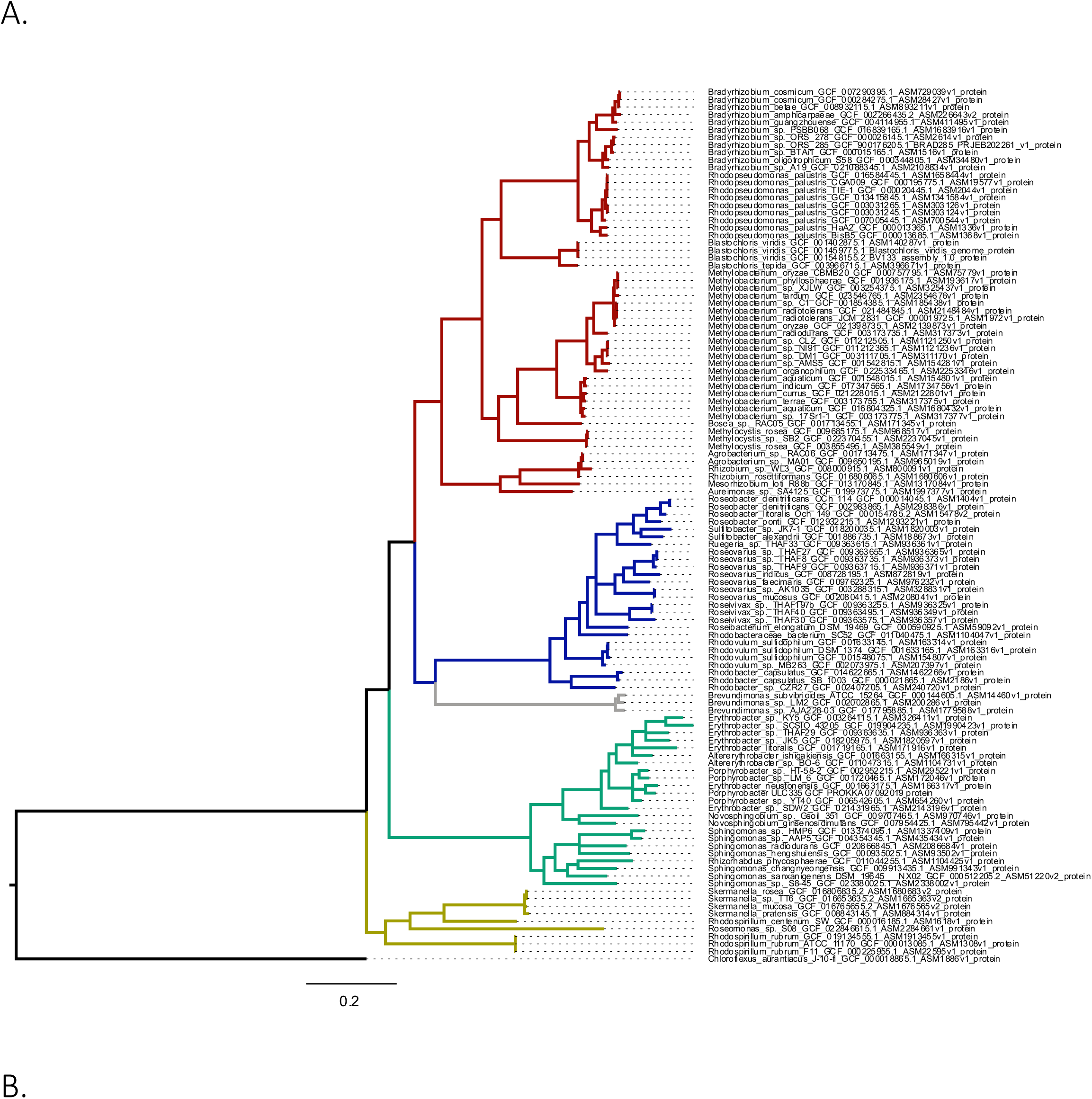
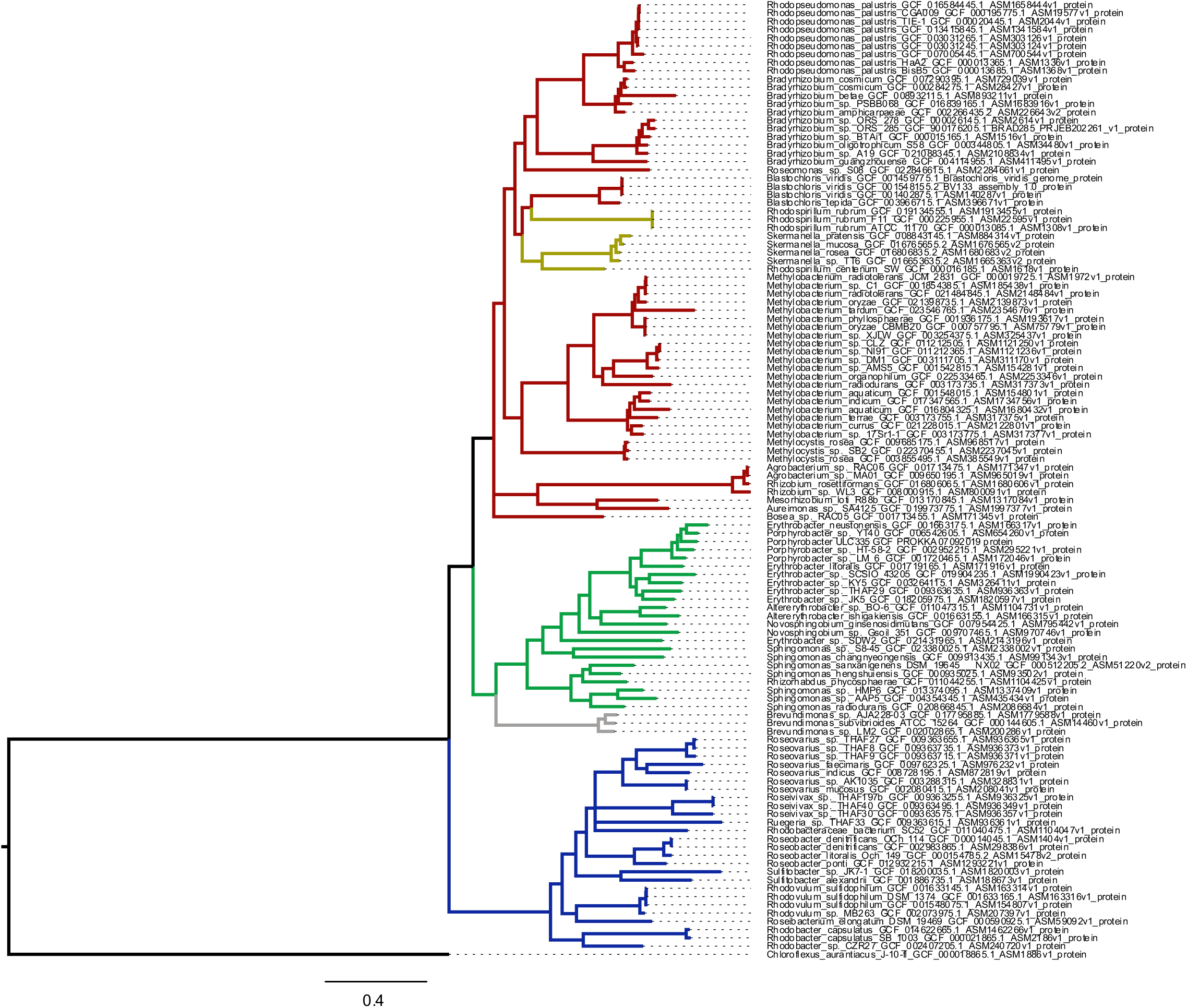
The anoxygenic phototrophy superoperon is primarily transferred vertically in alphaproteobacteria from the class level down. In general, the branching order is congruent, within classes, between a tree made from concatenated conserved core-genome proteins (A) and a tree of concatenated proteins belonging to the anoxygenic phototrophy operon (B). Exceptions are most likely due to horizontal gene transfer. Yellow: Rhodospirillales; green: Sphingomonadales; grey: Caulobacterales; blue: Rhodobacterales; red: Rhizobiales. Scale bar: average number of substitutions per site.

**SFigure 7:**
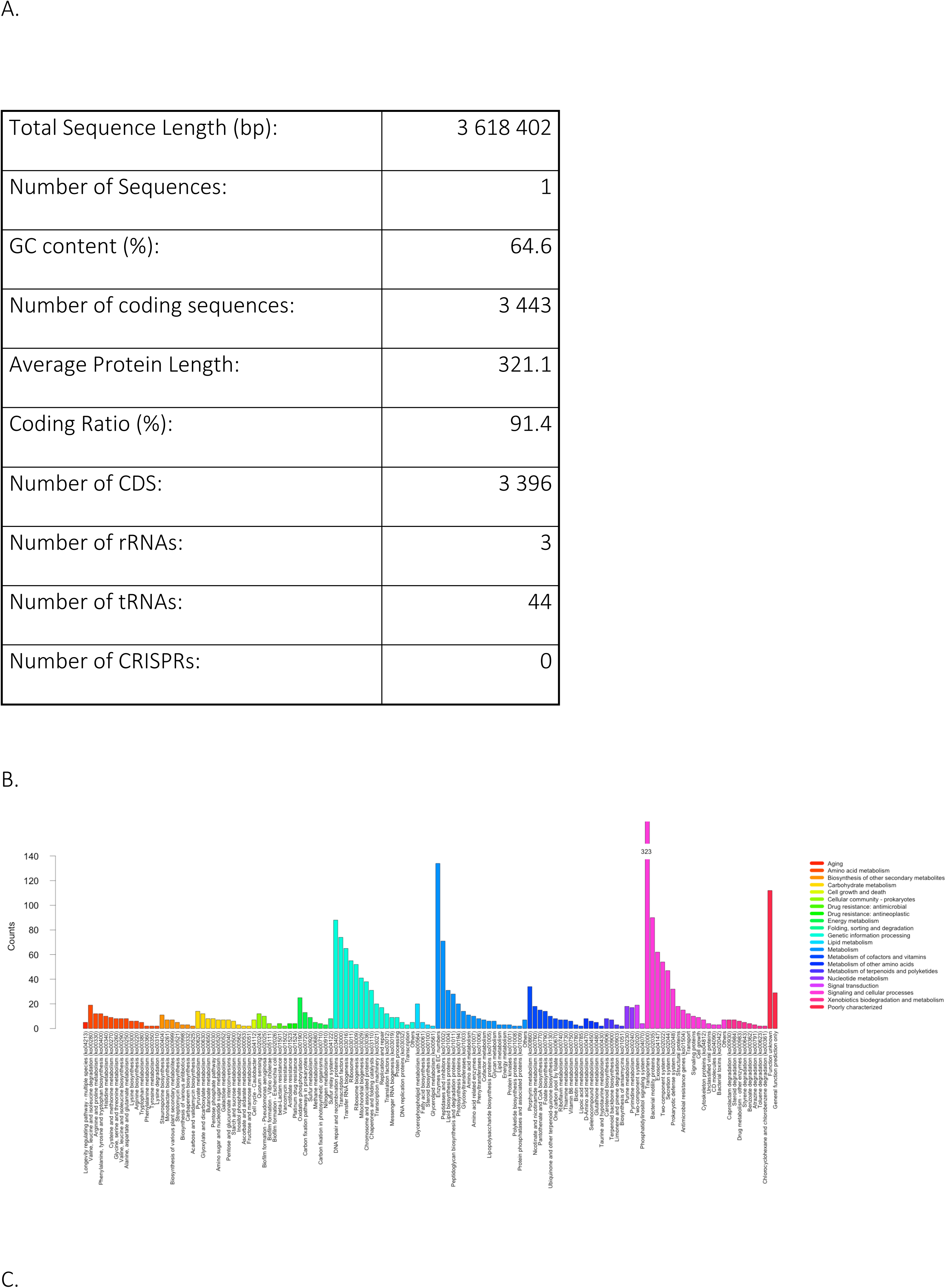
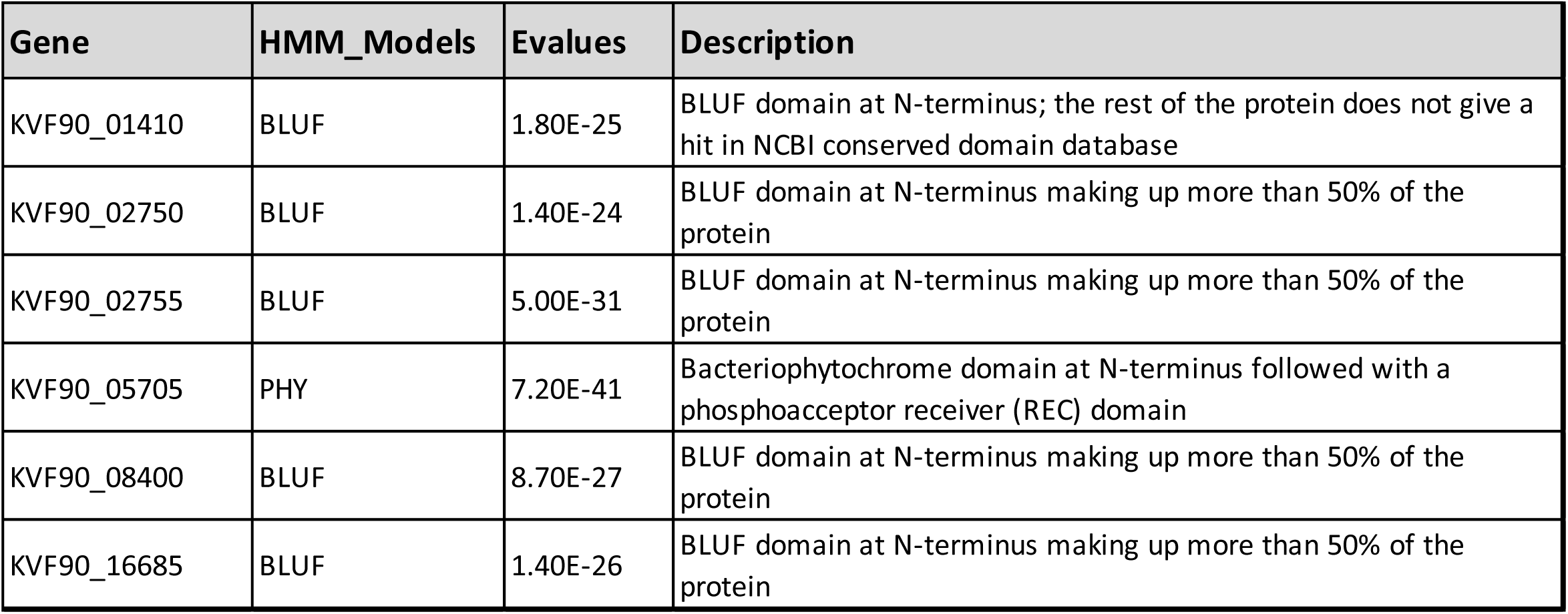
Global statistics and KEGG analysis of the genome of *Porphyrobacter* sp. ULC335. A. General characteristics of the genome of *Porphyrobacter* sp. ULC335. B. Functional classification of proteins encoded in the genome based on KEGG categories (two-levels). C. Table of all *Porphyrobacter* sp. ULC335 proteins containing light sensing domains (blue-sensing LOV or BLUF domains, or red-sensing bacteriophytochrome).

**SFigure 8:**
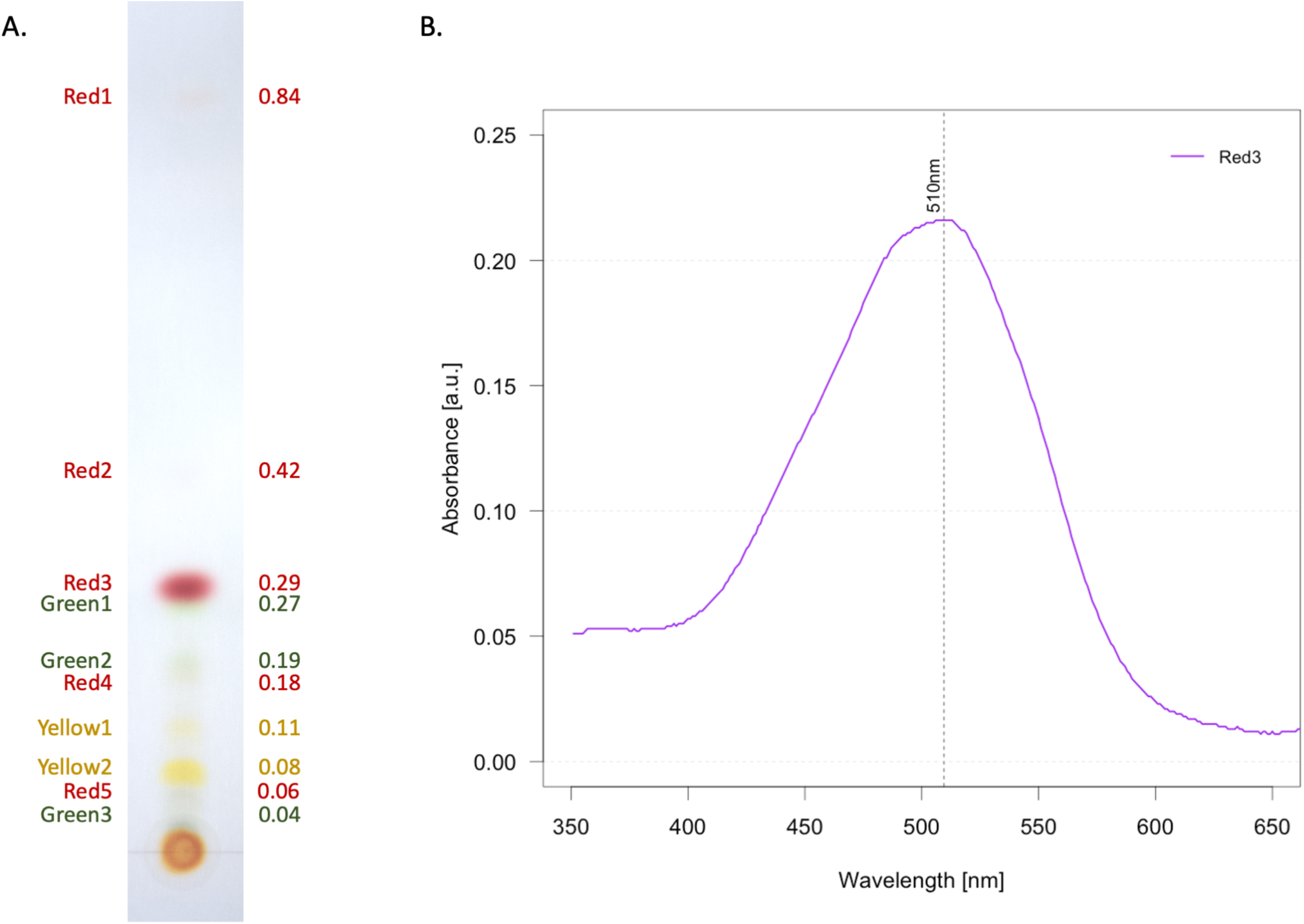
*Porphyrobacter* sp. ULC335 produces a range of carotenoids and bacteriochlorophyll *a* derivatives. A. Thin Layer Chromatography of pigments extracted from *Porphyrobacter* sp. ULC335. Red and yellow pigments are most likely carotenoids, green pigments precursors or derivatives of bacteriochlorophyll *a.* B. Absorbance spectrum of the main red pigment (“Red3”) eluted from the TLC plate. The UV-visible absorbance spectrum of the main purple-red cartotenoid is indicative of bacteriorubixanthinal, found in *Erythrobacter longus* and close relatives as the main reaction center-bound carotenoid[1–3].

**SFigure 9:**
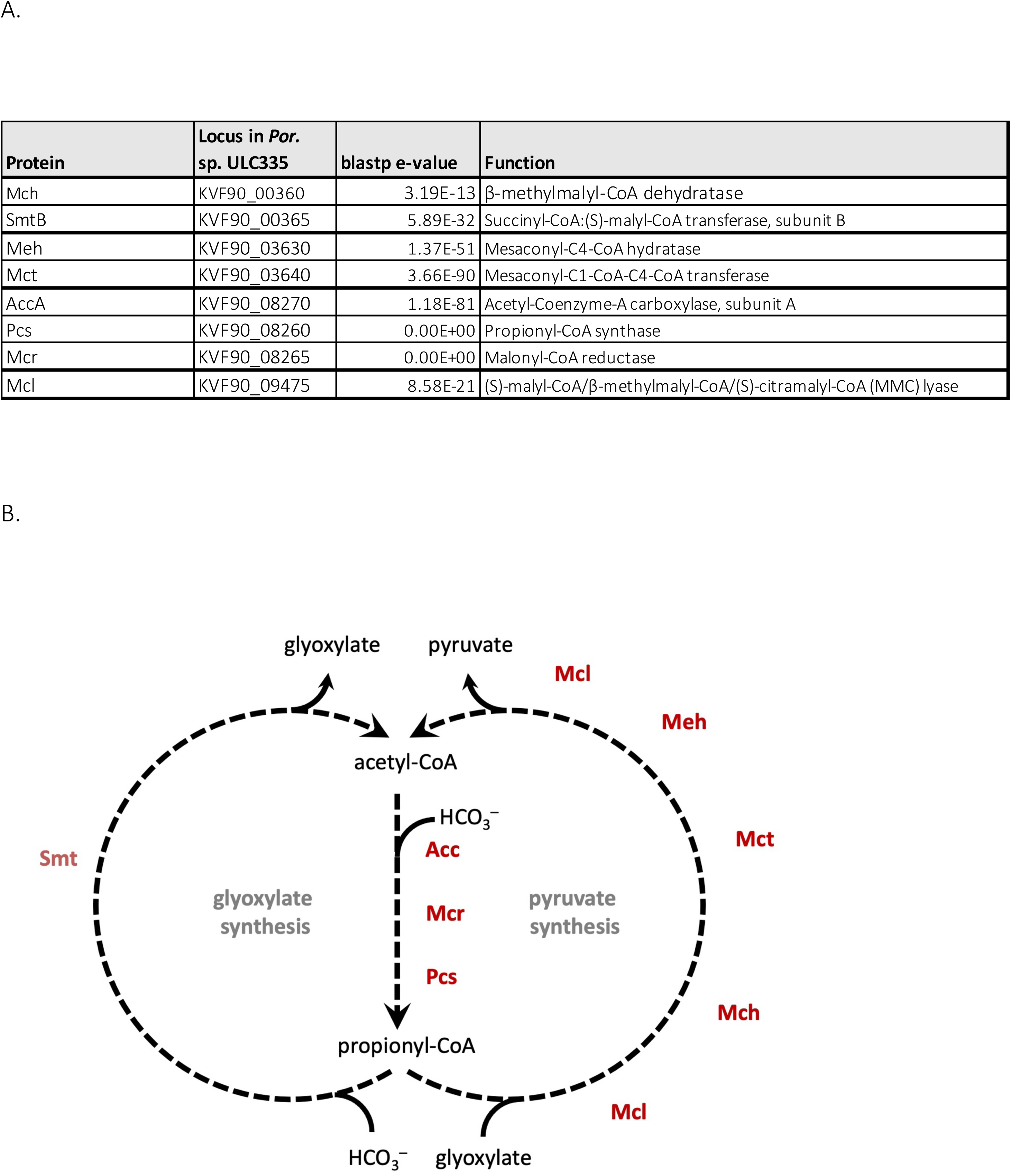
Homologs of several *Chloroflexus aurantiacus* enzymes participating to the 3hydroxypropionate bicycle for carbon fixation are found in *Porphyrobacter* sp. ULC335 genome. A. Table of enzymes found in the *Porphyrobacter* sp. ULC335 genome. The AccA protein is not considered a signature protein, but is often found associated with Pcs and Mcr homologs in genomes encoding the full 3-hydroxypropionate bicycle. The Mcl and the SmtB have relatively high evalues; an SmtA homolog is missing; the pathway is therefore considered incomplete like the one found in *Erythrobacter* sp. NAP1[4]. B. Schematic representation of the 3¬hydroxypropionate bicycle with the enzymes found the *Porphyrobacter* sp. ULC335 genome indicated in red.

**SFigure 10:**
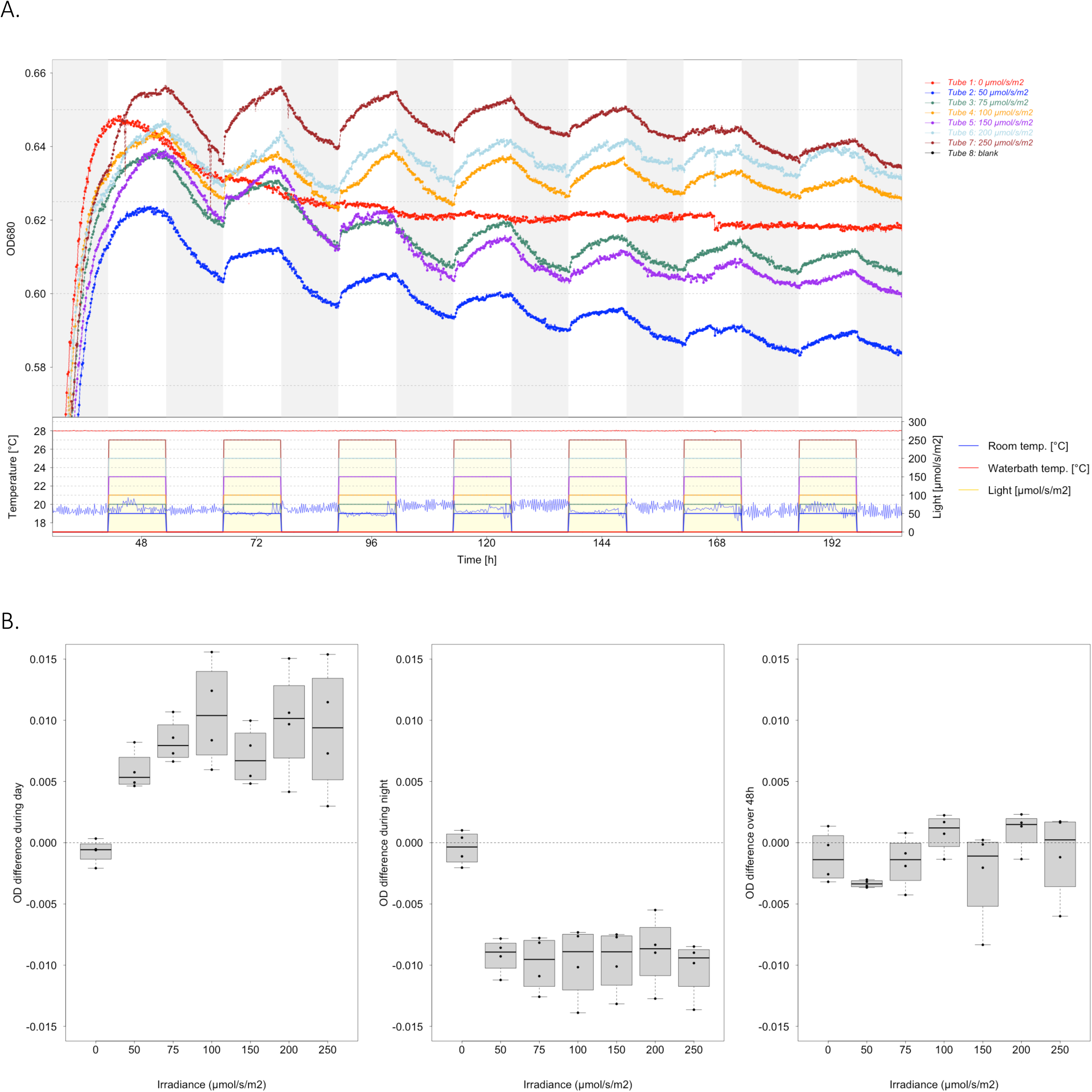
Growth curves of *Porphyrobacter* sp. ULC335 under 12:12 dark-light alternance in a range of light intensities. A. Rhythms are observed from 50 µmol/s/m2 to 250 µmol/s/m2; the maximal optical density tends to increase with light intensity. Growth curve above and conditions (light regime and water bath temperature; room temperature has no impact on culture conditions, but is added as a control) below. B. Boxplot showing the optical density differences (over the day, over the night and the net difference over 24 hours). The differences were calculated over four full cycles starting at 89h. The increase in density during the day correlates with the light intensity up to 100 µmol/s/m^2^, where it saturates; the decrease in density during the night seems dependent on light exposure but independent on light intensity; the net difference is close to zero except for the 50 µmol/s/m^2^ treatment, where the detrimental effect of light exposure observed during the night is not compensated by growth during the day.

**SFigure 11:**
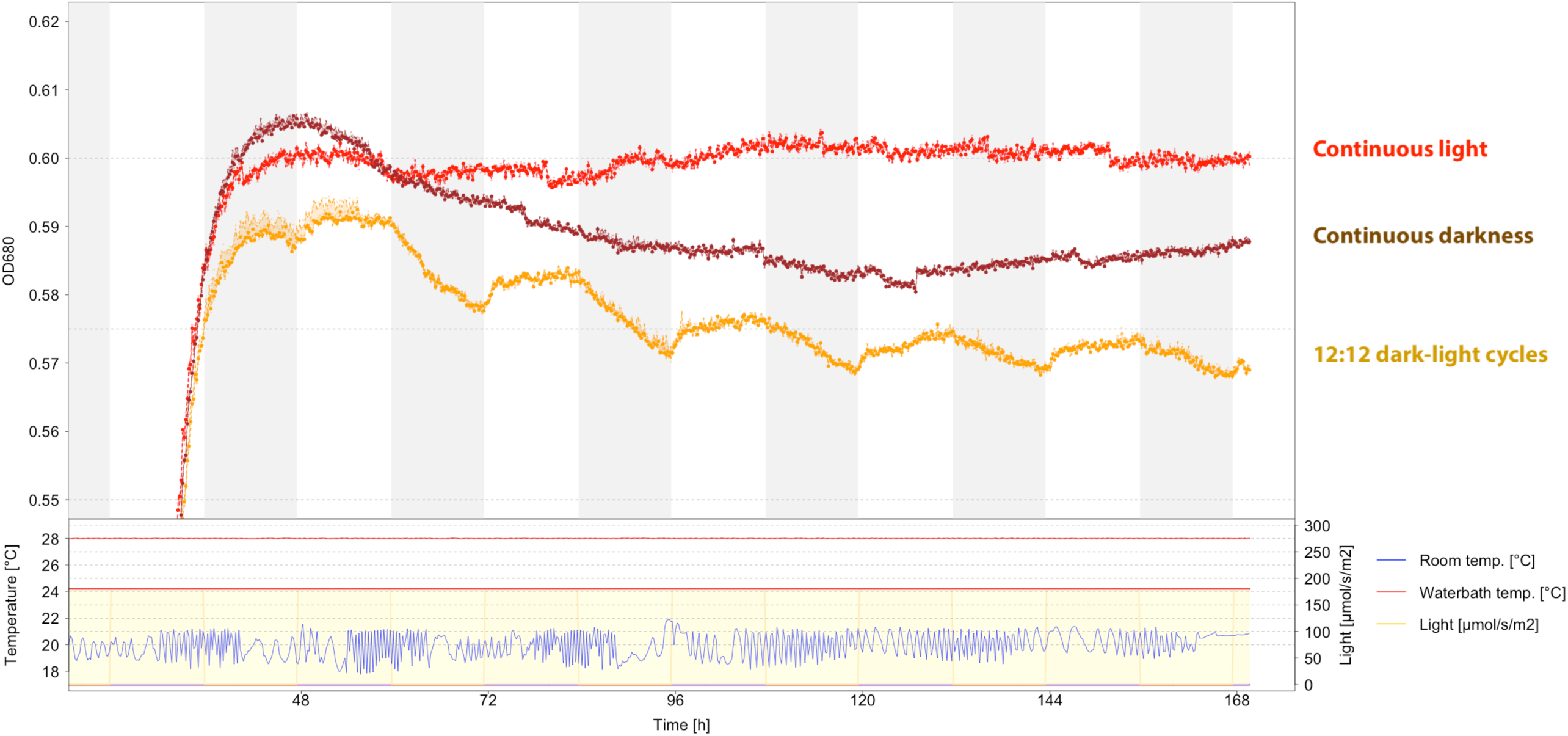
Growth curves of *Porphyrobacter* sp. ULC335 under continuous light (red), continuous darkness (dark brown), and 12:12 dark-light alternance (yellow), cultured in parallel. Rhythms are only observed under 12:12 dark-light alternance; the optical density under continuous light is consistently higher than under continuous darkness. Growth curve above and conditions (light regime and waterbath temperature; room temperature has no impact on culture conditions, but is added as a control) below.

**SFigure 12:**
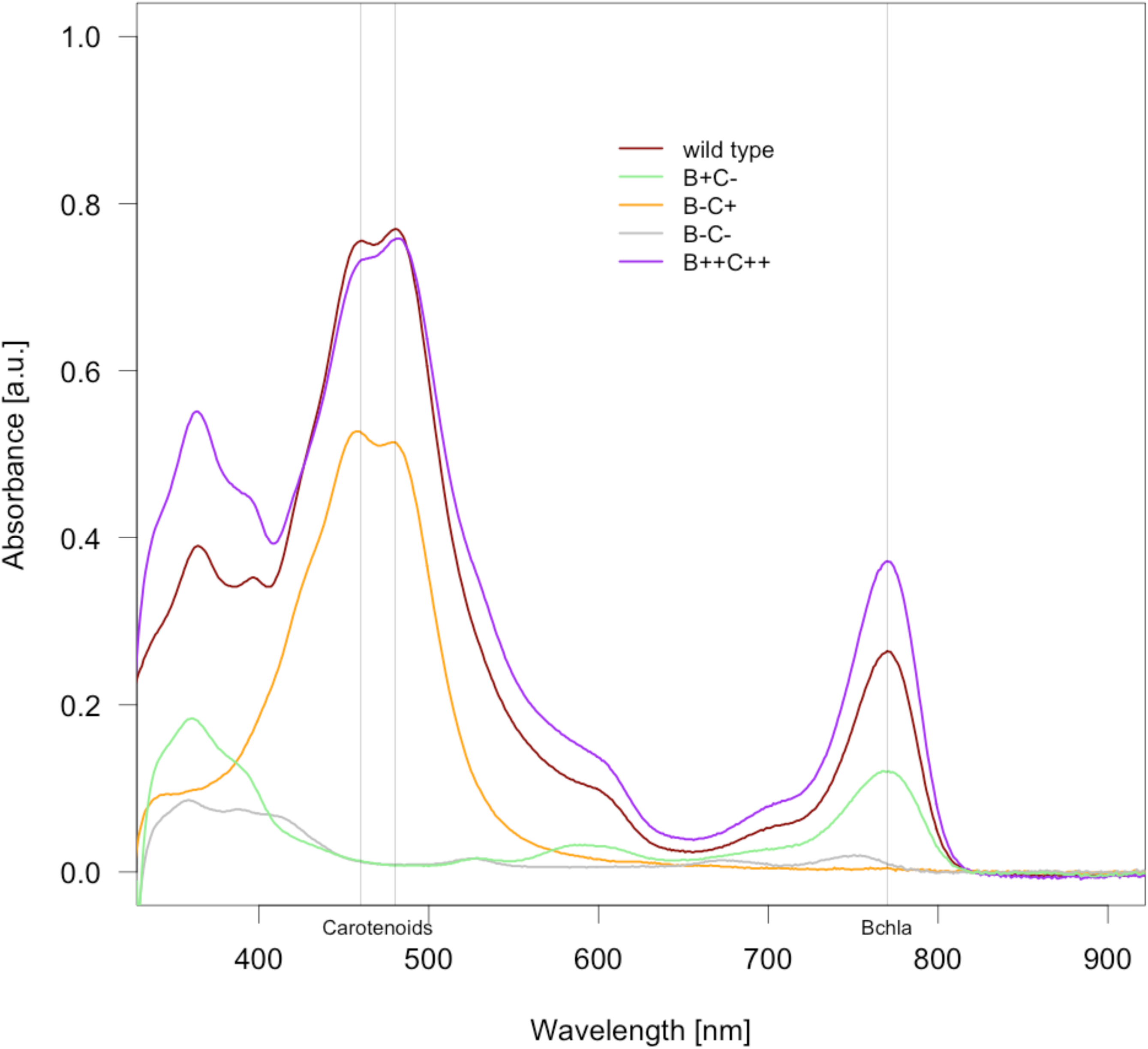
Absorbance spectra of *Porphyrobacter* sp. ULC335 and its four transposon mutants. The figure shows the absorbance profile of 7:2 acetone-methanol extracts of 72-hour batch cultures of *Porphyrobacter* sp. ULC335 and four mutants: the carotenoid mutant B^+^C^-^, the bacteriochlorophyll *a*-null mutant B^-^C^+^, which produces less carotenoids as well, the carotenoid-bacteriochlorophyll *a*-null mutant B^-^C^-^, and the carotenoid-bacteriochlorophyll *a* overproducer B^++^C^++^. Bacteria were grown in the dark at 30°C.

**SFigure 13:**
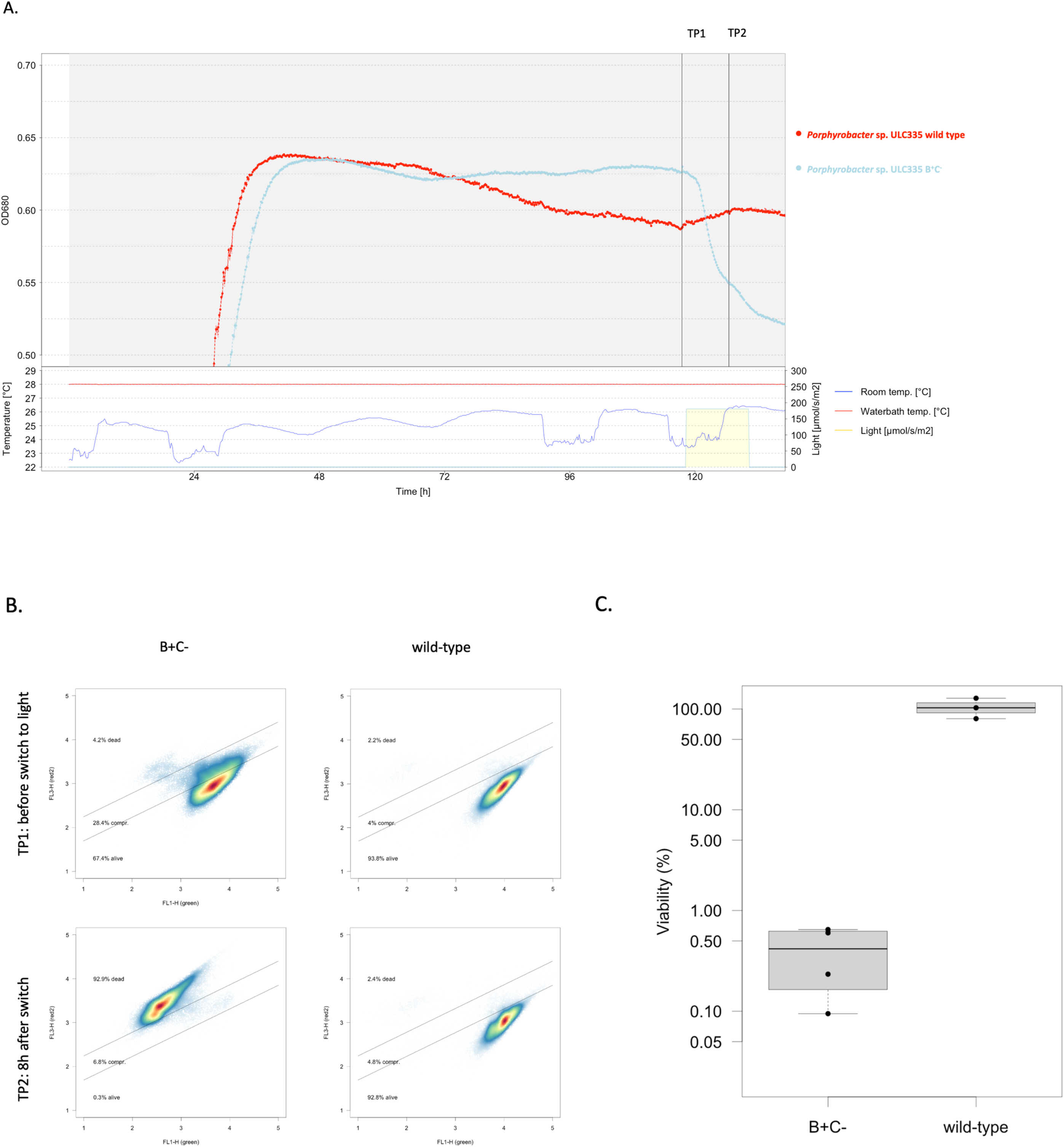
Upon light exposure strain B+C- loses viability. A. Cultures of *Porphyrobacter* sp. ULC335 wild type and the carotenoid mutant B+C- were grown in the dark for five days and then exposed to light (180 µmol/s/m2) for twelve hours. The growth curve shows a clear decrease in optical density right after exposure to light in the carotenoid mutant. B. The cultures were tested for viability right before and 8 hours after the switch to light using Life/Dead staining. The density graphs show that light induces cell permeabilization (death) in the B+C- strain but not in the wild type strain under the same conditions. C. Colony forming units [CFUs] confirm that the viability of the B+C- strain decreases by more than two orders of magnitude when the cultures are exposed to light while the viability of the wild type strain remains stable. The viability is the ratio of CFUs at TP1 divided by the CFUs at TP2 for each culture, expressed in percent.

**SFigure 14:**
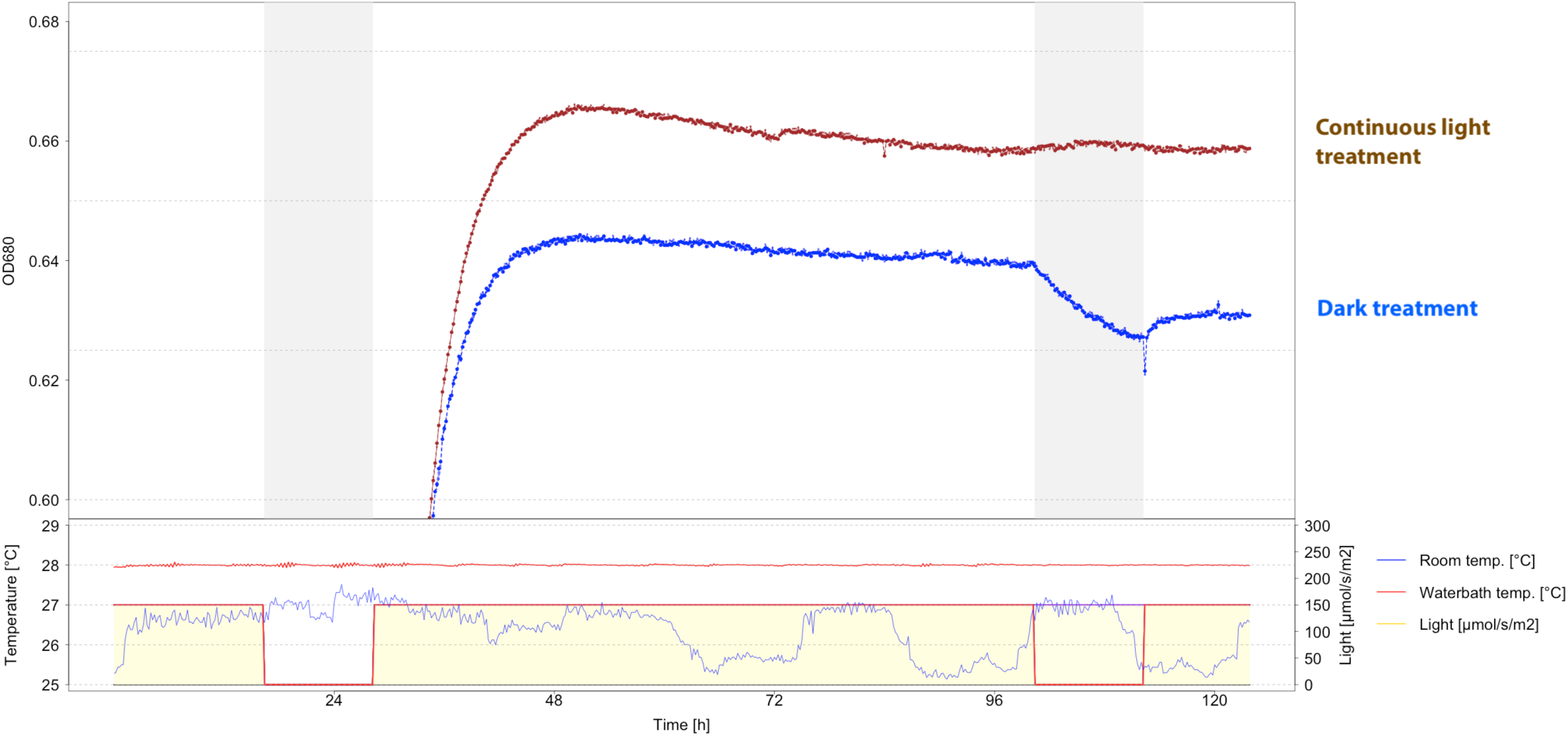
Actual data on which experimental design figure 3A is based. Growth curve above and conditions (light regime and waterbath temperature; room temperature is only indicative) below. The brown line is from a culture which has been maintained under continuous light as of stationary phase; the blue line is from a culture which has been switched to darkness at approximately 98 hours and switched back to light 12 hours after.

**SFigure 15:**
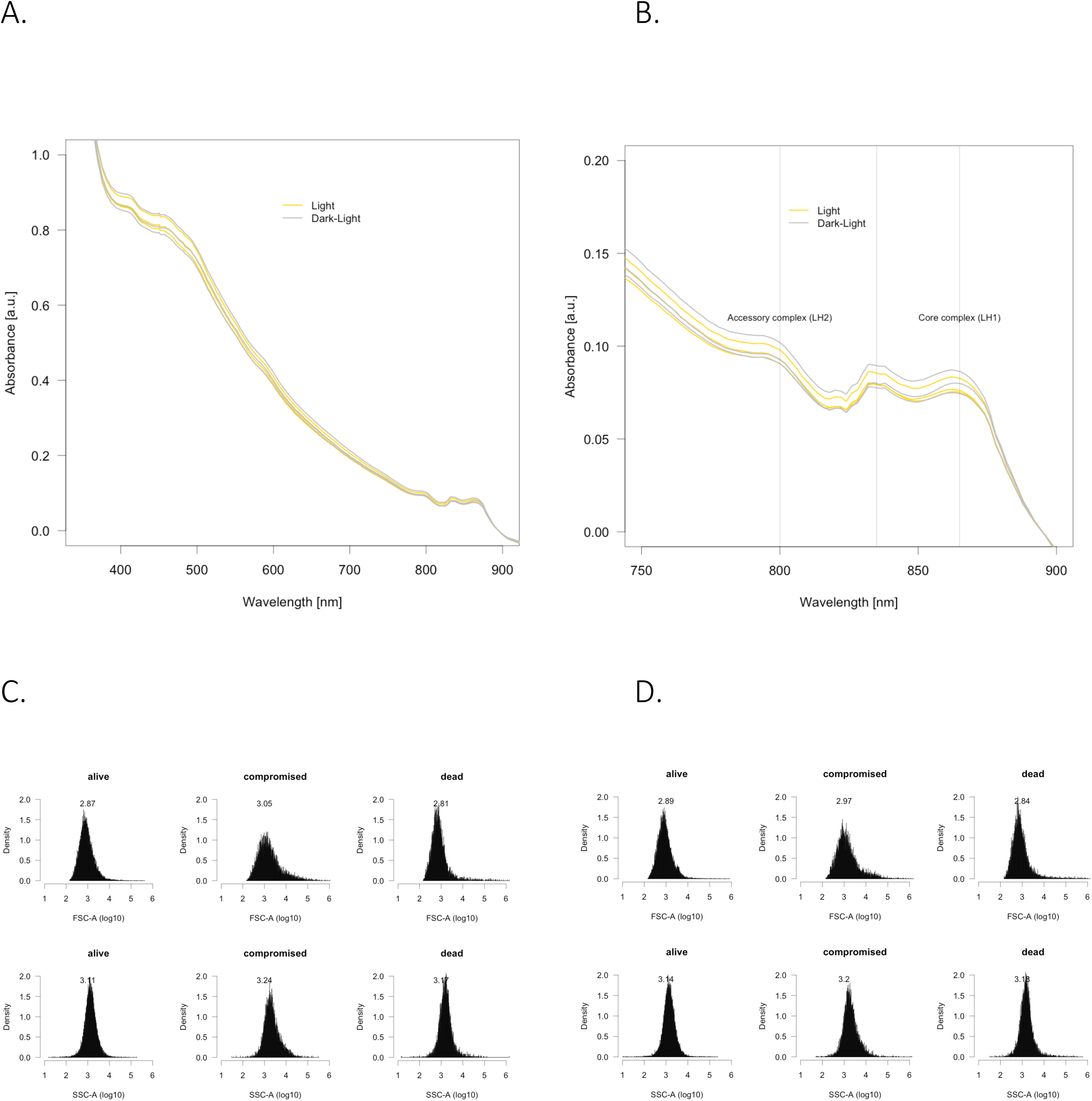
There is no difference in pigment content, cell size or cell aggregation between cells collected during a light or a dark phase. A. and B. Whole-cell absorbance spectra of five-fold concentrated *Porphyrobacter* sp. ULC335 cultures grown under continuous light (“Light”) or right after a 12-hour exposure to darkness (“Dark-Light”). A. Spectra from 350 to 900 nm showing carotenoid peaks between 450 and 520 nm and the light harvesting complexes between 750 and 900 nm. B. Detailed spectra from 750 to 900 nm showing the peaks corresponding to light harvesting complexes LH1 (800 nm) and LH2 (865 nm). The peak at 835 nm is atypical but could correspond to a secondary peak of the accessory complex (observed around 850nm in the anaerobic anoxygenic phototroph *Rhodobacter*)[5]. C. and D. Cell size (FSC-A) and cell granularity (SSC-A) of single cells from *Porphyrobacter* sp. ULC335 cultures grown under continuous light (C) or after a 12-hour exposure to darkness D); the values are ventilated based on their Live/Dead staining status. Neither the maxima nor the distributions show obvious differences between the two treatments suggesting that the cells are not different in size and do not aggregate in different ways in one treatment compared to the other

**SFigure 16:**
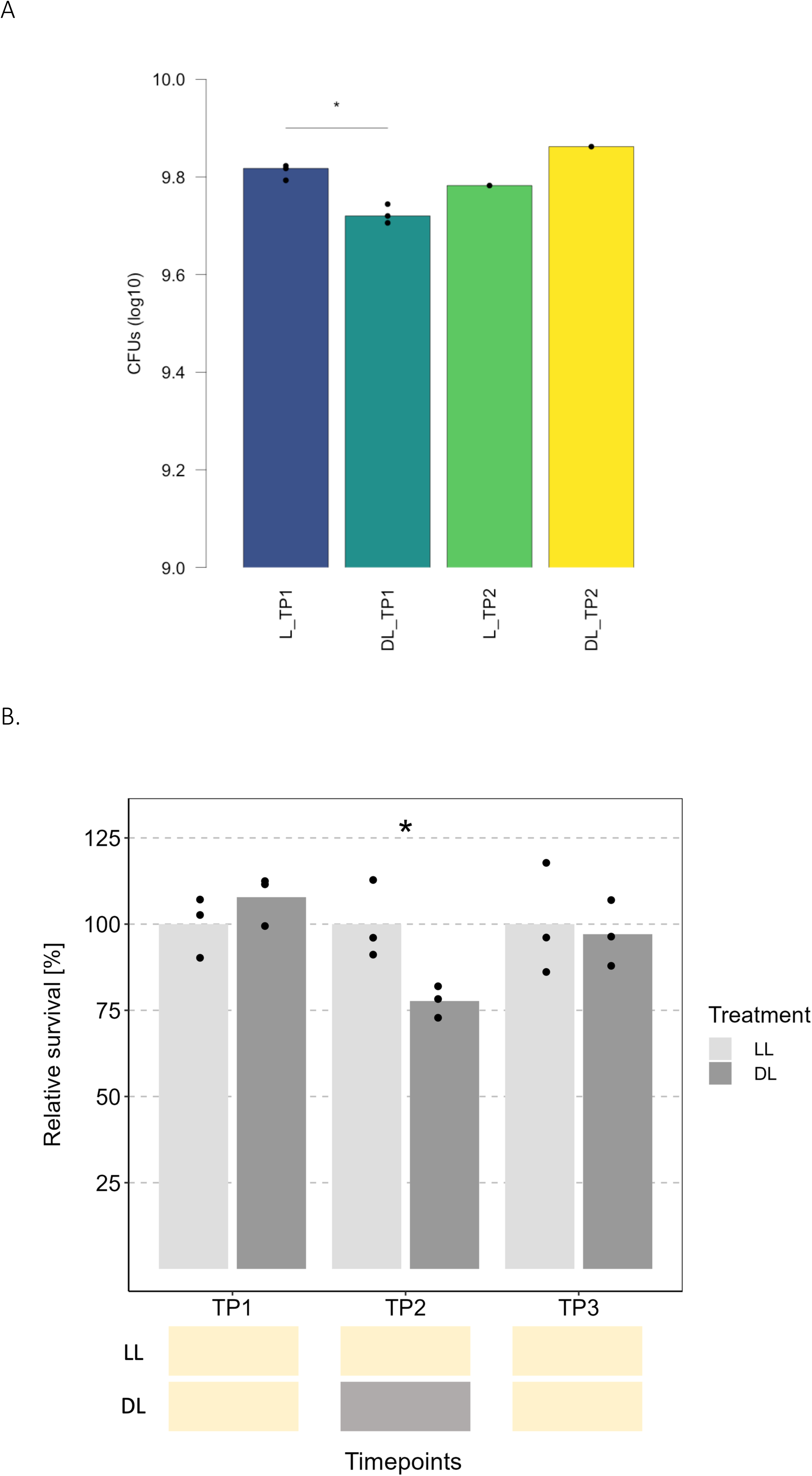
Colony forming units (CFUs) confirm the Live/Dead staining trend. A. This graph shows that the CFUs are significantly lower after an exposure to darkness than under continuous light. Only one sample was taken at the second time point, but it also confirms the recovery after reexposure to light. The star indicates a statistically significant difference (bilateral Student’s t.test, p-value<0.05). B. Results from an independent experiment, which show exactly the same trend. Six *Porphyrobacter* sp. ULC335 independent cultures were grown to stationary phase and maintained under continuous light (150 µmol/s/m^2^) for 72 hours; to allow for Bchl*a* accumulation, cultures were exposed to darkness for 12 hours during exponential phase. Then, for three replicates (LDL treatment), light was switched off for 12 hours and switched back on, while the three other tubes were maintained under continuous light (LL treatment). Samples were taken from all six cultures one hour before the switch to darkness, six hours after switch to darkness, and twelve hours after re-exposure to light. In the DL treatment, the viability decreases significantly (paired bilateral Student’s t.test, p-value<0.05) during the period of darkness compared to the control and increases back after re-exposure to light. Star: p-value < 0.05.

**SFigure 17:**
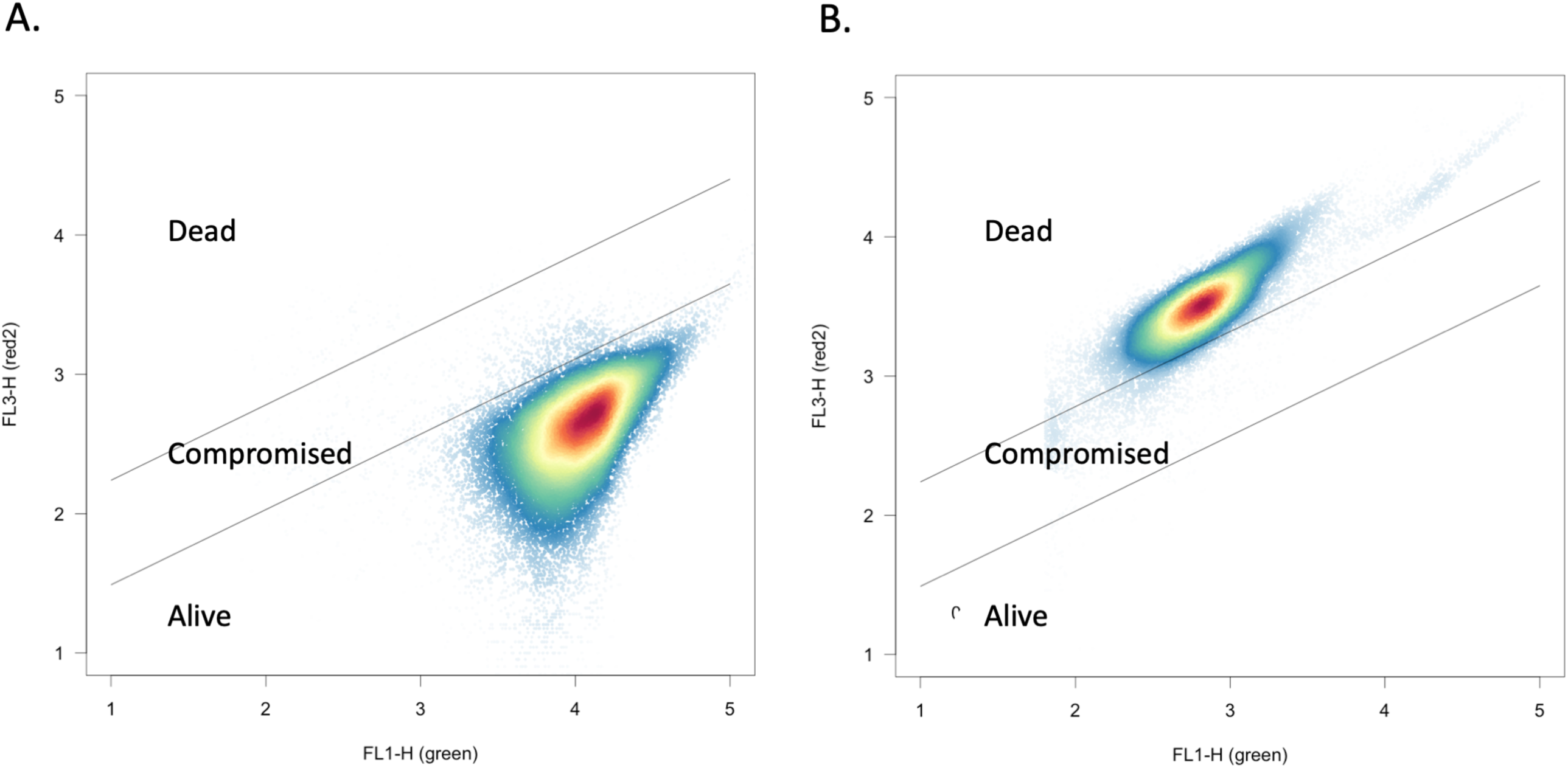
Distribution of green and red fluorescence values in live and dead *Porphyrobacter* sp. ULC335 cells. A. Live cells from a 36-hour culture show high green (FL-1) and low red signal (FL-3). B. Cells fixed with 2.5% glutaraldehyde show low green and high red signal. The zone in between the lines accommodate the high green and high red signal cells, that we consider “compromised” and in which the “dead dye”, propidium iodide, can enter, but at a slow speed.

**SFigure 18:**
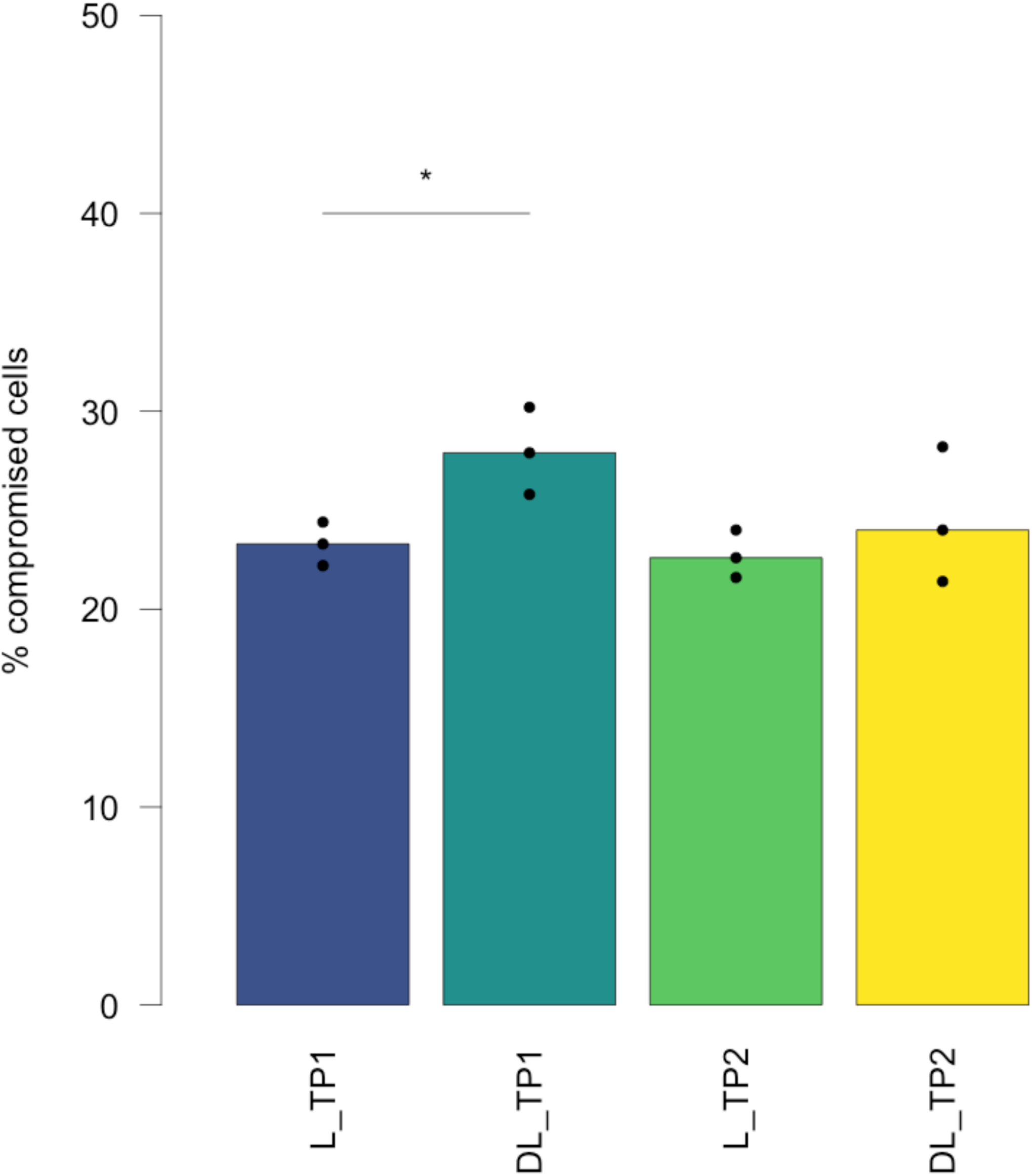
The proportion of compromised cells is greater in the cultures subjected to a dark phase than in the cultures maintained under continuous light. This graph shows the proportion of cells falling in the “compromised” category, based on flow cytometry, is significantly higher after an exposure to 12-hour darkness than under continuous light. Like for dead cells, the proportions go back to baseline. The star indicates a statistically significant difference (bilateral Student’s t.test, p- value<0.05).

**SFigure 19:**
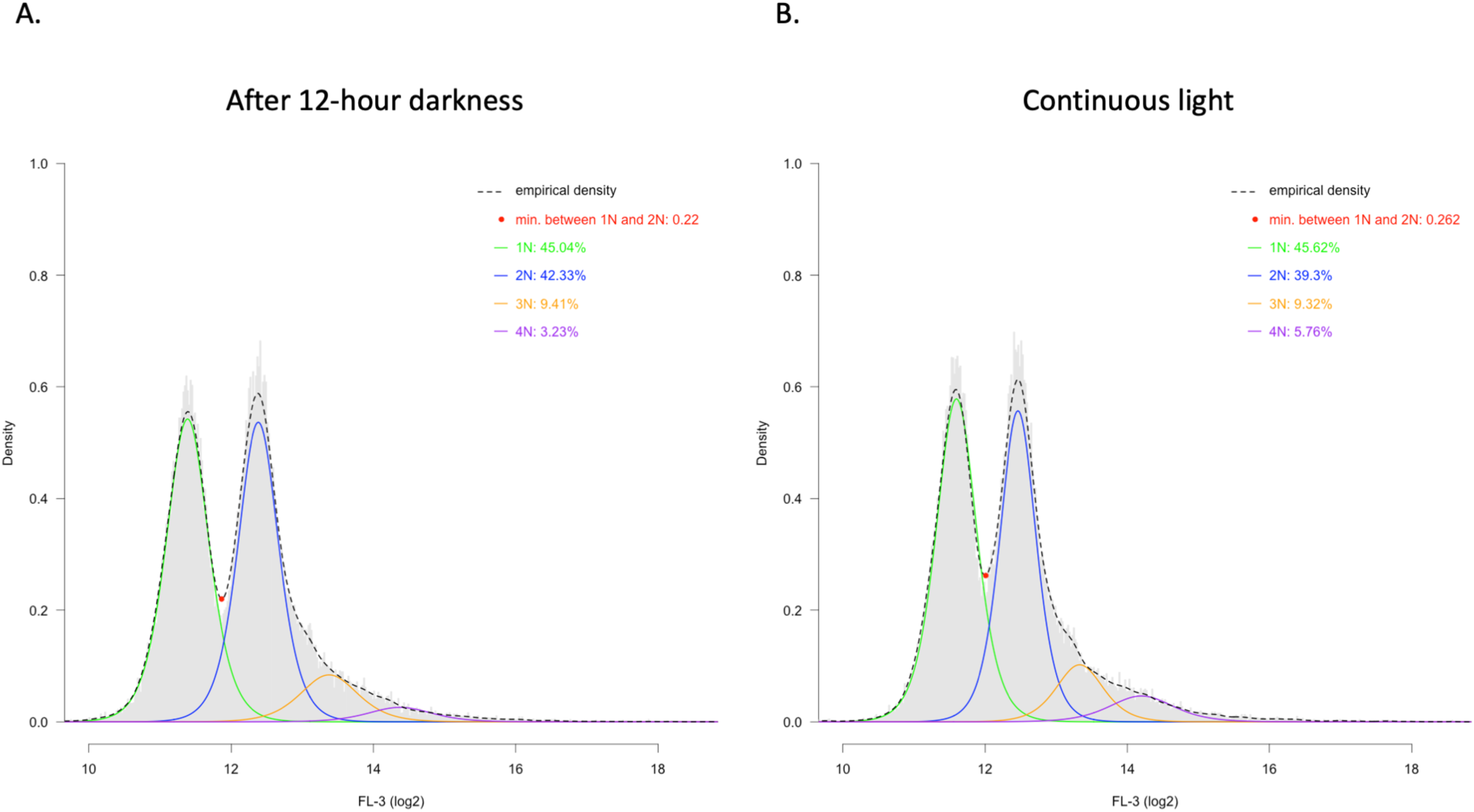
The replicating-cell population size differs between the dark (A) and continuous light (B) treatments. The graphs show a histogram of the distribution of green fluorescence (FL-1) in DyeCycle Orange-stained cells as measured by flow cytometry. Overlapping the histogram, lines represent the smoothed empirical density over the population (ticked black line), the modelled 1N (green line), 2N (blue line), 3N (orange line), and 4N (purple line) chromosome content peaks as predicted from the location of the 1N and 2N peaks (local maxima and distance between two N values). In both treatments, more than 85% of the cells were likely arrested at 1N-chromosome (before the start of DNA replication) or at 2N chromosomes (after the end of chromosomal replication and before cell division) The relative proportions between the four peak were not different between treatments. However the lowest empirical density value between the 1N and the 2N peaks, which is a measure of the actively replicating population, was significantly lower in cells exposed to the dark than in the cells maintained under continuous light.

**SFigure 20:**
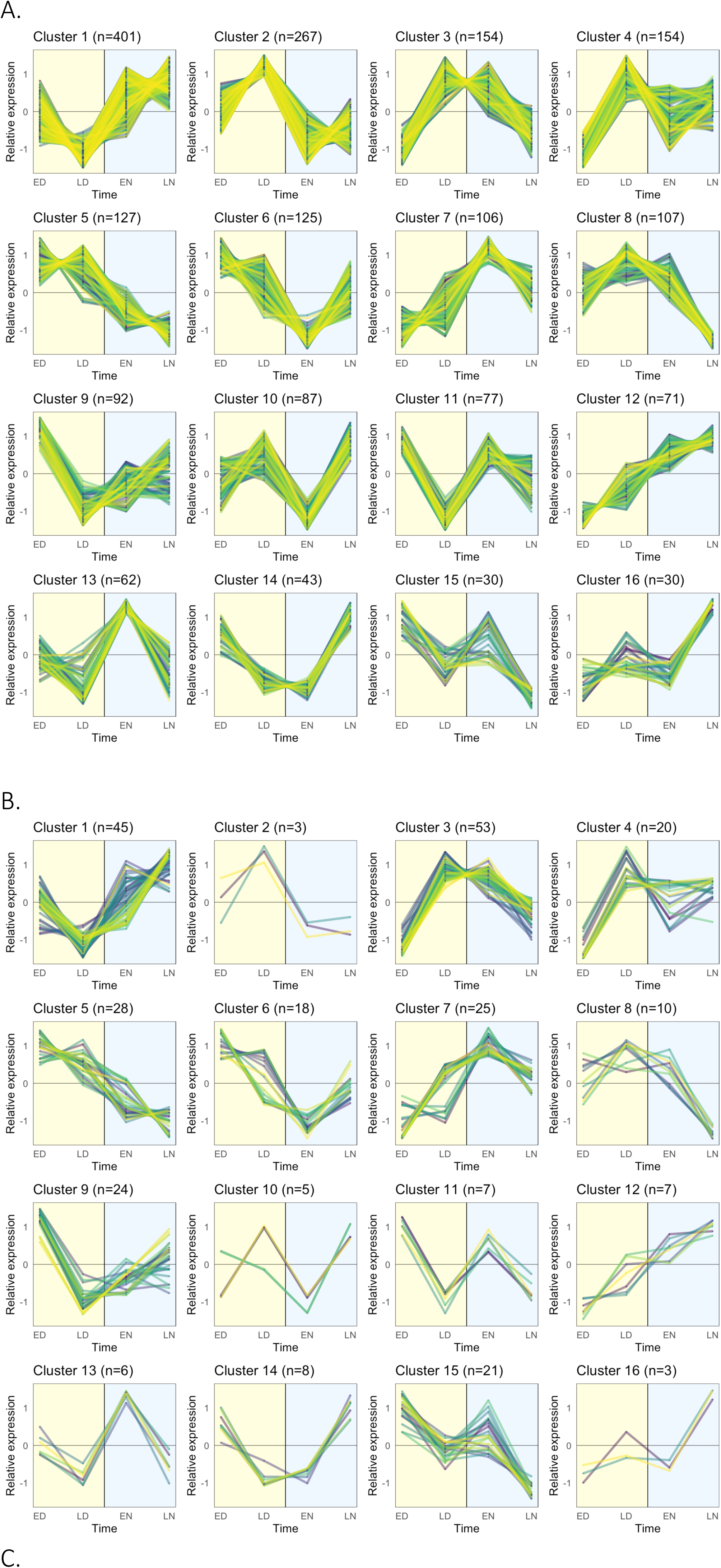
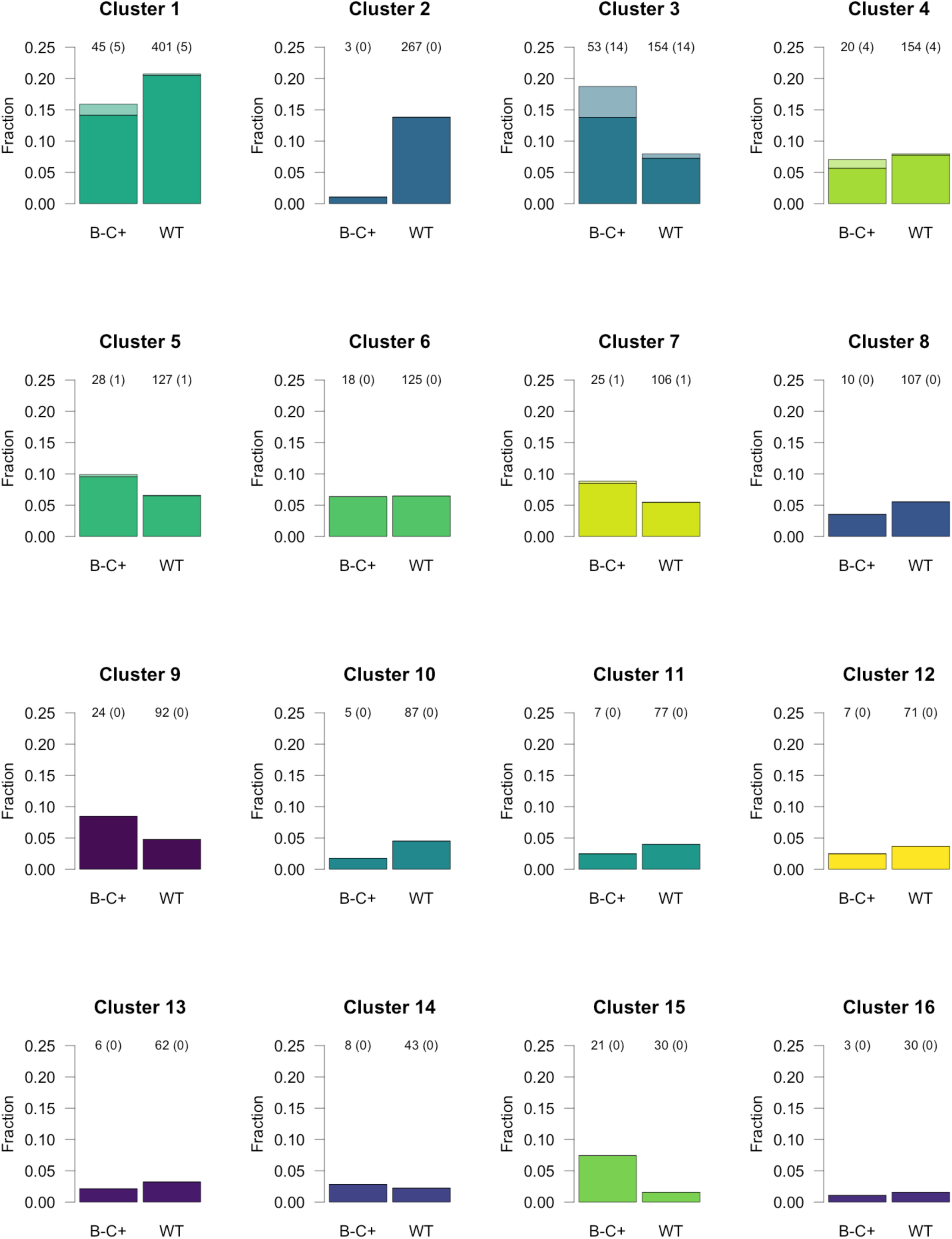
Clusters of temporally regulated genes in the wild type (A) and the B^-^ C^+^ mutant (B), and their representation in each strain (C). The figure summarizes information about the temporal regulation of genes in the wild type strain and its Bchl*a* null mutant. A. and B. The clusters of genes showing a consistent light-response pattern in either the wild type (A) or the photosynthesis null mutant B^-^C^+^ (B) are presented based on their temporal profile. The datapoints represent the relative transcription rates, i.e. the Z-scores of the rlog-transformed count data]. C. The barplots shows the fraction each cluster represents among temporally regulated genes in either (dark colors) or both (light colors) strains.

**SFigure 21:**
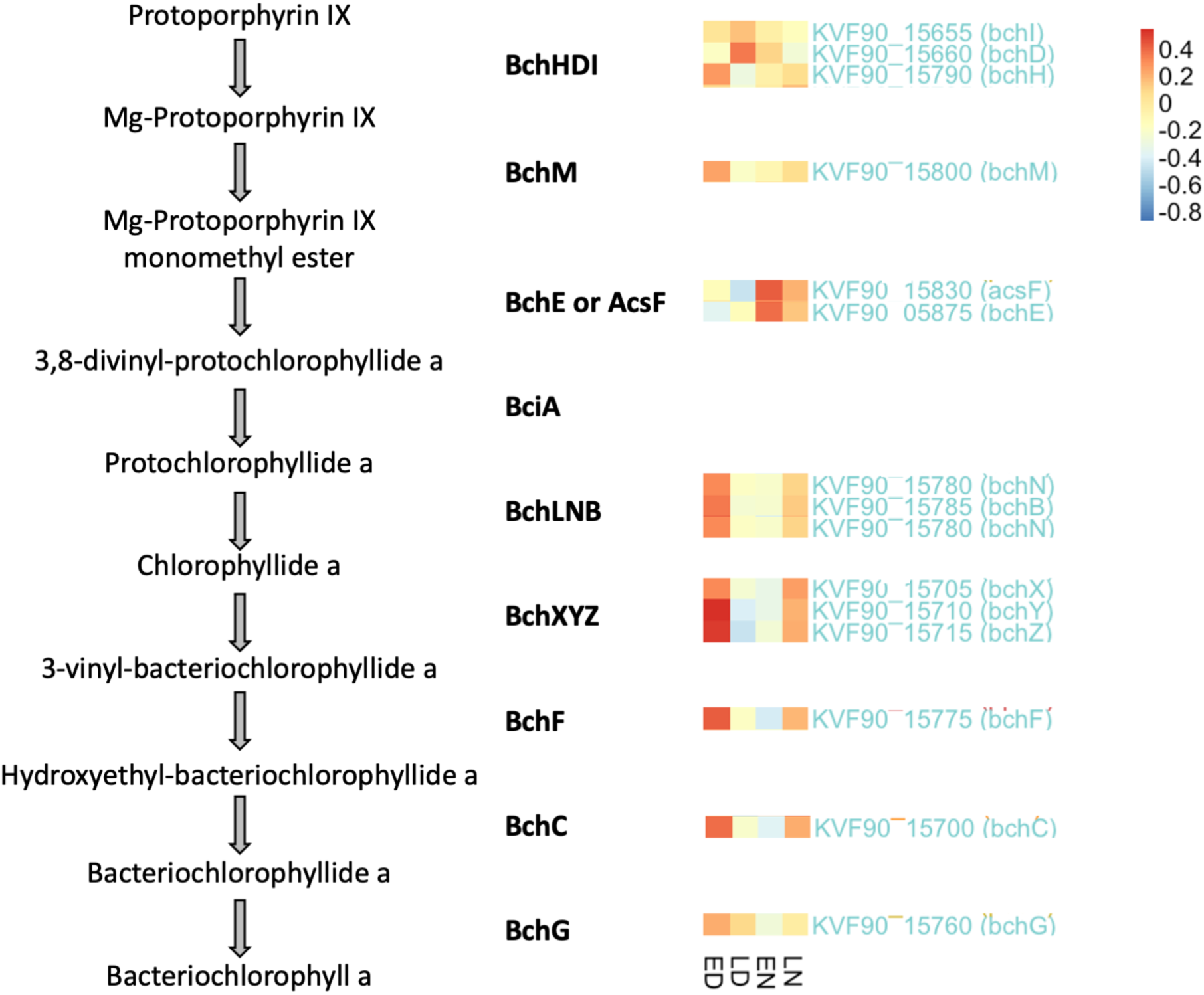
Schematic representation of the bacteriochlorophyll *a* biosynthesis pathway together with the enzymes involved and their transcriptional profile in the wild type strain. The heatmaps have been normalized to the wild type strain. Three temporal profiles emerge: 1. *bchDI,* maximally expressed at LD, 2*. bchHM, bchLNB, bchXYZ, bchF* and *bchC*, expressed at LN and ED, 3. *bchE* and *acsF*, expressed at EN and LN.

**SFigure 22:**
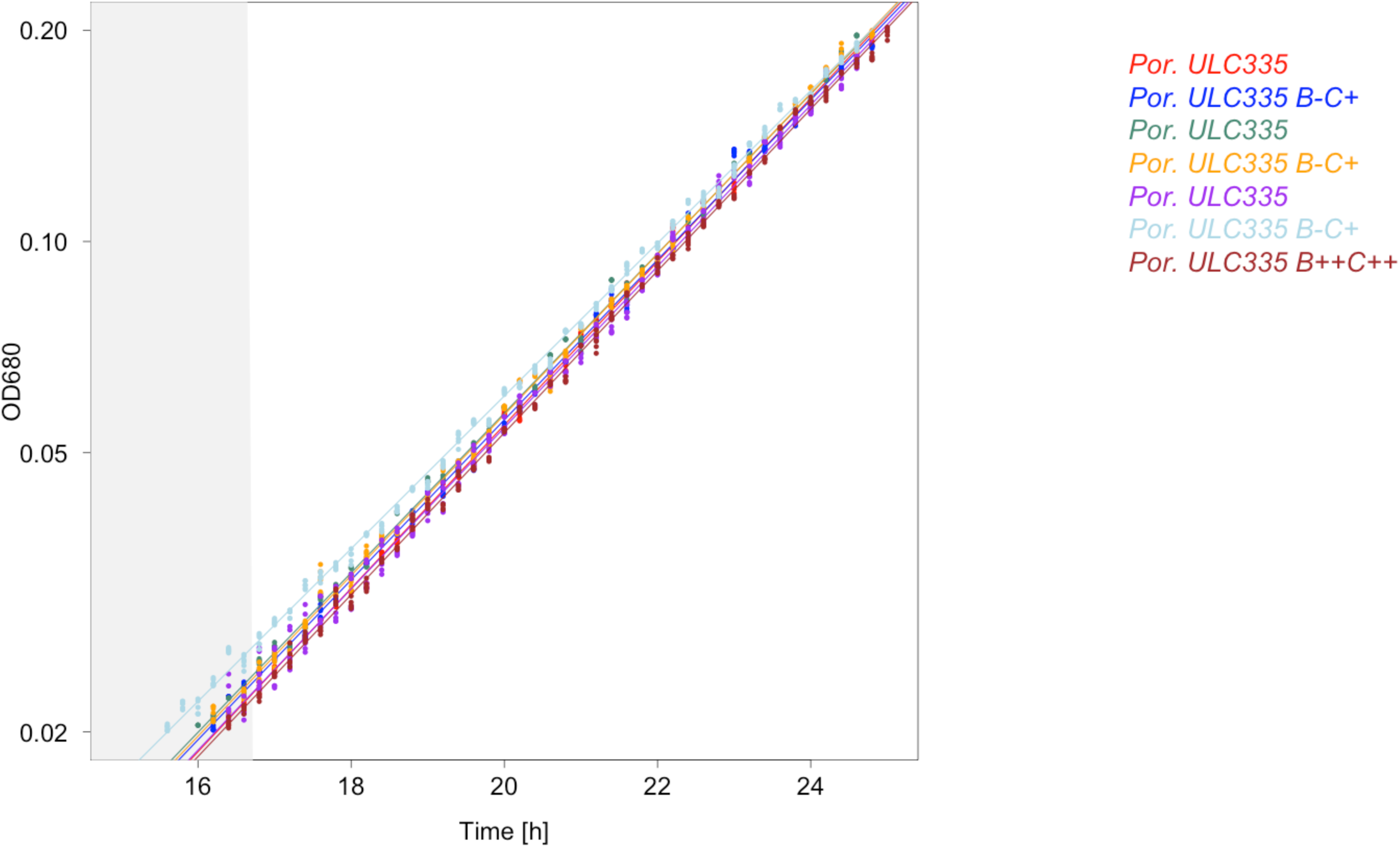
The growth rates of the wild type, the Bchl*a*-null B^-^C^+^ and the Bchl*a*-carotenoid overproducer B^++^C++ strains do not differ significantly. The figure represents the OD680 as a function of time of three wild type, three B^-^C^+^, and one B^++^C^++^ replicates in early exponential phase. The slope does not significantly differ between the cultures, indicating that the growth rate of the three strains in exponential phase is very similar.

**SFigure 23:**
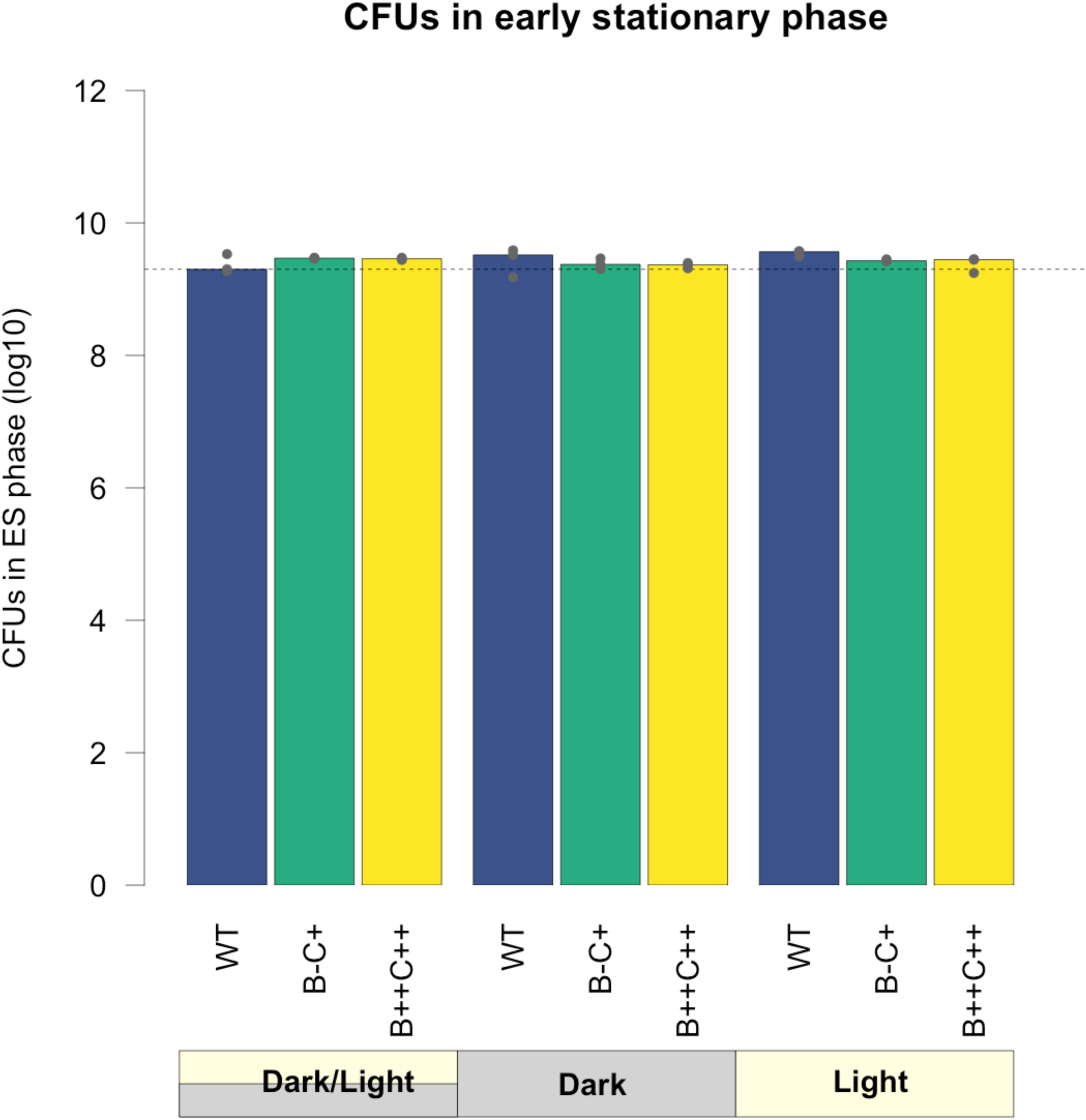
The colony forming units (CFUs) of the three strains do not differ significantly in early stationary phase (after 48 hours), whatever the illumination regime. Histograms of CFUs obtained through plating the initial timepoint of experiments shown in Figure 5B. The treatments were compared using ANOVA (p-value > 0.05).

**SFigure 24:**
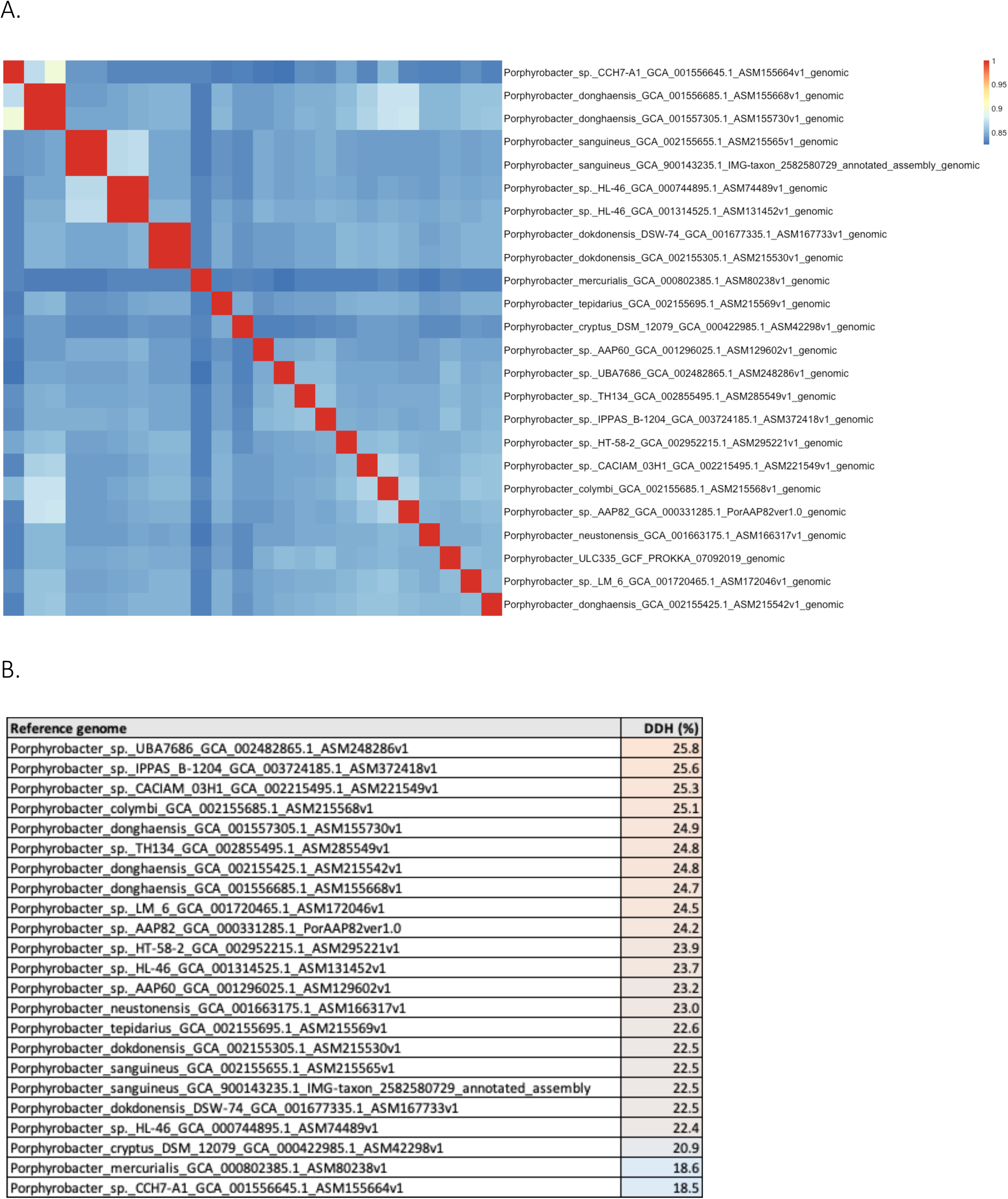
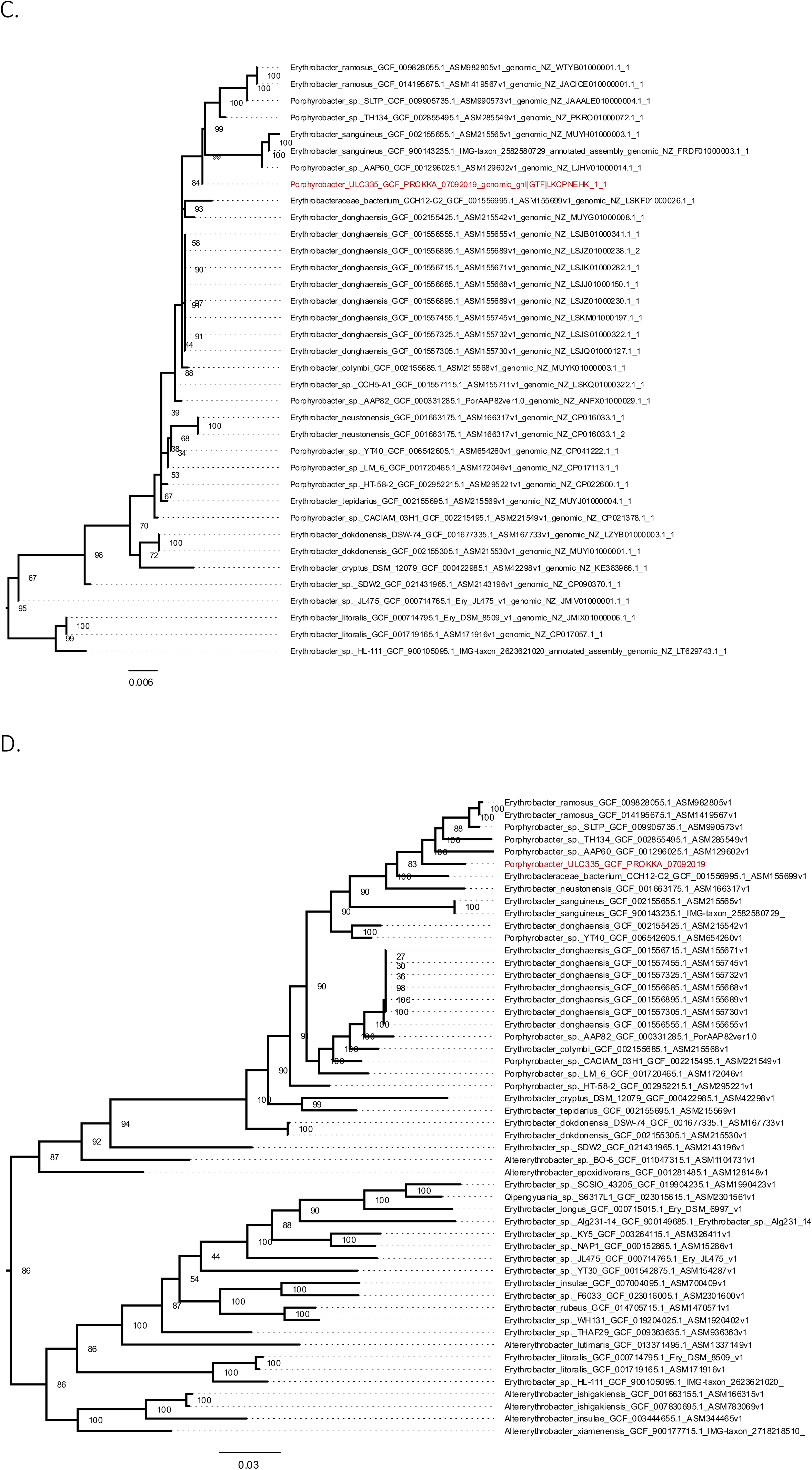
Average nucleotide identity [ANI] and digital DNA-DNA hybridization [DDH] between the genomes of *Porphyrobacter* sp. ULC335 and several other *Porphyrobacter* strains representative of the diversity of the genus. A. ANI as calculated using PYANI [6]. The ANI of strain ULC335 is lower than 0.85 for a representative set of *Porphyrobacter* strains or species, which suggests that strain ULC335 may be a new *Porphyrobacter* species. B. The DDH was estimated using the digital Genome-to-Genome Distance Calculator algorithm recommended by the Leibniz Institute DSMZ (Formula 2) [7]. The DDH between *Porphyrobacter* sp. ULC335 and a representative set of *Porphyrobacter* strains or species is estimated below 30%, which suggests that strain ULC335 may be a new *Porphyrobacter* species. C. Phylogenetic tree of the closest relatives of strain ULC335 based on the full 16S rRNA gene sequence. D. Phylogenetic tree of the closest relatives of strain ULC335 based on 27 concatenated conserved proteins. Strain ULC335 is shown in red. *iqtree* bootstrap values are shown at the nodes (max. 100). Scale bars: average number of substitutions per site.

## Supplementary Methods

### RESSOURCE AVAILABILITY

#### Lead contact

Requests for information or strains should be directed to the lead contact, Diego Gonzalez.

#### Materials availability

*Porphyrobacter* sp. ULC335 and its transposon mutants are available from the lead contact.

#### Data and code availability

The *Porphyrobacter* sp. ULC335 genome is publicly available on the NCBI repository (SAMN19986570). The RNA-sequencing data is publicly available on the NCBI Gene Expression Omnibus (GEO) repository (GEOXXX).

### EXPERIMENTAL MODEL AND SUBJECT DETAILS

*Porphyrobacter* sp. ULC335 was isolated from a BG11 Petri dish on which *Snowella* sp. ULC335 and its associated community of heterotrophs were plated. It formed isolated small orange to red colonies on the agar. After two initial reisolation steps on PYE [8], it was confirmed pure by microscopy and stocked at −80°C in 20% glycerol. Further cultivation was done on BG11-P (see below). The ULC335 community, the *Porphyrobacter* sp. ULC335 strain and its transposon mutants used in this article are listed and referenced in the table below.

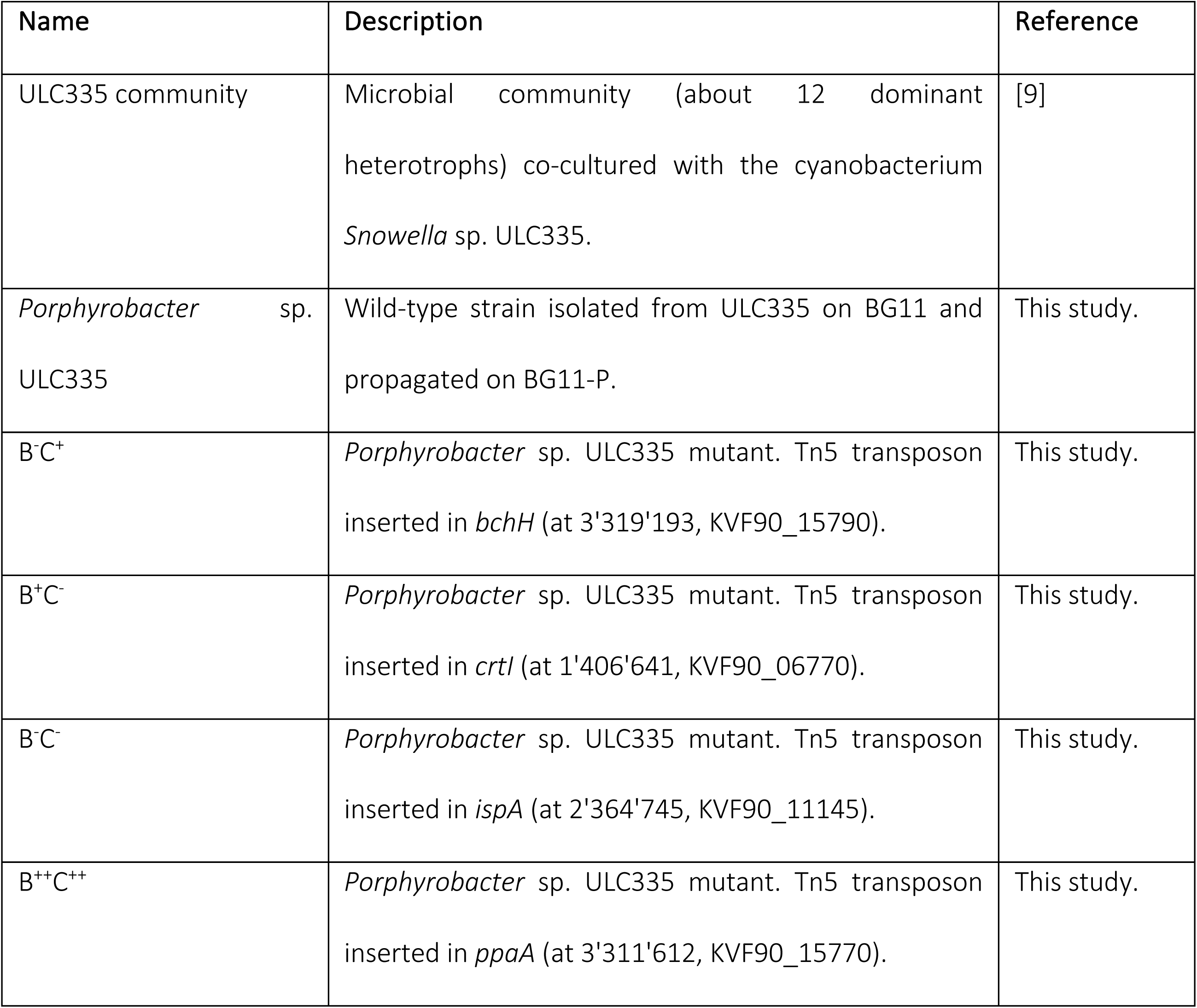

### METHOD DETAILS

#### Standard growth conditions

*Porphyrobacter* sp. ULC335 was grown in BG11 media [10] supplemented with 0.5g/L Tryptone (Oxoid) and 0.25g/L Yeast Extract (Oxoid), as well as 1/100 dilution of vitamin solution (hereafter, BG11-P). Vitamin solution contained (mg/L): Biotin (4), Folic acid (4), Pyridoxine-HCl (20), Riboflavine (10), Thiamine-HCl 2H_2_O (10), Nicotinamide (10), D-Ca-pantothenate (10), Vitamin B12 (0.2), p-Aminobenzoic acid (10). Standard batch growth took place at 30°C with agitation. Growth in the Photon Systems Instruments Multi-Cultivator MC1000 cultivating device took place at 28°C. Batch cultures for competition and survival experiments were done in 24-well plates at 28°C with agitation. *E. coli* was grown in LB (tryptone, 10 g/L; yeast extract 5 g/L; NaCl, 5 g/L; tryptone and yeast extract from Oxoid) at 37°C. Solid media were supplemented with 1.5% technical agar.

#### Real-time optical density monitoring

Real-time monitoring of the optical density of *Porphyrobacter* sp. ULC335 cultures was done in a Multi-cultivator MC1000 (Photon Systems Instruments) under warm-white LEDs at 150 to 180µmol/s/cm^2^ at 28°C with mixing ensured by bubbling (eight culture tubes in parallel). OD680 was measured every 12 minutes (six individual measures per timepoint) with one tube used as blank to correct for measurement fluctuations due to changes in the room temperature. Transitions in illumination regimes (square wave) are reported on the graphs presented for each experiment. Cells were grown in BG11-P from an initial optical density <0.001.

#### UV-visible-close IR spectra

SFigure 2. The cell pellets collected from 12 ml of 48-hour old liquid cultures of *P. neustonensis* DSM9434 and *Porphyrobacter* sp. ULC335 were extracted with 700 µl of 7:2 acetone-methanol solvent for 5 minutes; after centrifugation, the absorbance of the supernatant was measured over the UV-visible-close IR spectrum in quartz cuvettes (1 cm path length) with a UV-visible spectrophotometer (Thermo Scientific Genesys 10s).

SFigure 11. For whole-cell spectra, cells were washed once with NaCl 0.9% and resuspended five-fold concentrated in NaCl 0.9%. 350 to 900 nm spectra were obtained from 200µl in a 96-well plate with a spectrophotometer (SpectraMax i3x, Molecular Devices).

#### Thin Layer Chromatography

50 ml of three day-old batch cultures of relevant strains were centrifuged; the pellet was resuspended in 20 ml of NaCl 0.9% and centrifuged again. The pellet was resuspended into 1 ml of 7:2 acetone-methanol and left in the dark for 5 minutes; the supernatant was collected by centrifugation. About 100µl of supernatant were spotted onto a Silica 60 F_254_ Thin Layer Chromatography (TLC) plate (Merk, art. 5554), previously dehydrated at 120°C for one hour, and left to dry. The TLC plate was migrated in 8:0.75:0.2:0.8:0.25 petroleum ether:hexane:isopropanol:acetone: methanol. Migration front and spots were used to calculate retardation factors for each pigment. The main red pigment was eluted into 7:2 acetone-methanol and its UV-visible-close IR spectrum was measured in quartz cuvettes (1 cm path length) as stated above.

#### Transposon library

5ml of 48-hours old *Porphyrobacter* sp. ULC335 culture were pelleted, washed twice with BG11-P, and combined with an equivalent pellet of a conjugation-competent *E. coli* strain harbouring pRL27 [11]. The mixture of bacteria was spotted on a BG11-P plate and incubated at 30°C for 4 hours. The bacterial spot was then collected and spread on a total of twenty BG11-P supplemented with 50µg/ml kanamycin and 10µg/ml chloramphenicol (to counterselect *E. coli*). About 5’000 colonies were collected and stocked at −80°C.

A collection of mutants whose color differed from the wild type was isolated on BG11-P plates or BG11 plates supplemented with 2.25g/L Tryptone and 0.75g/L Yeast extract. The B^+^C^-^ and B^-^C^-^ strains were white; the B^-^C^+^ strain was orange to light red rather than dark red like the wild type; the B^++^C^++^ was darker than the wild type. The mutants were restreaked twice to make sure that they were pure. The insertion sites were identified using the method previously published [11]. Briefly, the genomic DNA of the mutants was extracted using an ammonium-acetated-based protein precipitation protocol followed by ethanol-precipitation [12]. 1 µg of purified DNA was digested using BamHI (1 hour, 37°C); after deactivation of the restriction enzyme, the DNA was autoligated over night at 4°C using the T4 ligase (NEB). The ligation product was transformed into *E. coli* DH5a-λpir using the heat shock method. The plasmid was extracted and the insert was sequenced using the tpnRL17–1 primer (Microsynth). The insertion sites were identified using *blast* [13].

#### pufM PCR

SFigure 1. The *pufM* PCR was carried out with pufM-F and pufM-R primers using GoTaq (M3001, Promega) on genomic DNA extracted using Quick-DNA Fungal/Bacterial Miniprep Kit (Zymo Research) from 500µl of over-night liquid culture in BG11-P. Annealing temperature was 55°C.

#### Genome sequencing and annotation

The genomic DNA was purified from 3ml of *Porphyrobacter* sp. ULC335 48-hour culture using the Qiagen Genomic-tip 20/g kit. Sequencing was performed using the Pacific Biotechnology technology (PacBio Sequel). The sequencer generated 38’568 polymerase reads (mean length: 31,681) and 262’709 subreads (mean length: 4’389 bps); mean final coverage was 314. The genome was assembled using PacBio CCS 4.1.0 for the read correction step and Flye 2.6 [14] with a target size of 5MB, 1000bps minimal overlap between reads, and five final polishing steps for the assembly. In its final state, the genome comprises only one circular contig.

The functional classification of the genome content was done using the online blastKOALA service (https://www.kegg.jp/blastkoala/) with the genus *Porphyrobacter* as a reference. The two first levels of the KEGG classification were used to make SFigure 7.

#### RNA-sequencing

The wildtype and B^-^C^+^ strains were cultured under 12h/12h dark-light alternance with constant oxygenation in the MC1000 cultivation device (warm white LEDs, 150 µmol/s/cm^2^). After three days in stationary phase, samples were taken at four different time points from three different replicates. Sampling was done one hour before and one hour after the light-dark or dark-light transition: Early Day (ED) was taken one hour after dark-light transition, Late Day (LD) one hour before light-dark transition, Early Night (EN) one hour after light-dark transition, Late Night (LN) one hour before dark-light transition. RNA was extracted from 1 ml culture (OD650=0.3) using TRIzol (Invitrogen) followed by purification on columns. Briefly, cells were lysed in 1 ml TRIzol Reagent at 65°C for 10 minutes and kept at room temperature for 5 minutes; 200 µl of chloroform were added and, after mixing by inversion, incubation for 3 minutes at room temperature and centrifugation (15 minutes, 12000 x g, 4° C), 500 µl of the upper phase were mixed with an equal volume of 70% EtOH and loaded on a RNeasy mini-kit column (Qiagen). After an on-column DNase treatment (RNase-free DNase set, Qiagen), columns were washed and RNA eluted in 40 µl of RNase-free H2O according to the manual instructions. RNA was quantified using Nanodrop2000 (ThermoFisher Scientific) and Qubit 2.0 fluorometer (Qubit RNA Broad Range Assay kit, ThermoFisher Scientific), checked for degradation on an agarose gel, and checked for possible DNA contamination by qPCR using SybrGreen (Rotor-Gene SYBR Green PCR kit, Qiagen) on a Rotor-Gene Q thermocycler (Qiagen). RNA-sequencing was carried out at Novogene on Illumina NovaSeq systems after ribosomal depletion and in-house library construction. An average of 40 to 70 million 150 bps pair-end reads were obtained per sample.

Reads were trimmed and filtered using *fastP* [15], and then mapped on the *Porphyrobacter* sp. ULC335 genome using *bowtie2* [16]. The number of reads per coding sequence was counted using *featureCounts* [17]. Exploratory analysis and statistical testing were carried out using *R*. A first analysis, based on the *DESeq2* package [18] and workflow, was used to determine which genes were differentially expressed in the wildtype and B^-^C^+^ strains at each of the four time points (pairwise comparisons for each timepoint, adjusted p-value < 0.01). A second analysis, based on the *ImpulseDE2* package [19], was used to determine which genes were temporally regulated across the four time points in either the wild-type or the mutant strain (adjusted p-value < 0.01). To assign each temporally regulated gene to a specific temporal profile (cluster), we used *hclust* (Euclidian algorithm, 15 clusters) on the Z-scores of the *rlog*-transformed count data for all four time points. The *ggVennDiagram* [20] and *ggplot* [21] libraries were used to plot the summary figures.

#### Phylogenetic analysis

The proteomes of alphaproteobacterial genera containing at least four complete genomic sequences were downloaded from the NCBI repository and searched, using *blastp* [13] (evalue<10E-5), for homologs of fourteen proteins belonging to the anoxygenic phototrophy superoperon (BchH, BchD, BchB, BchY, BchZ, BchN, PpsR, BchC, BchX, BchI, BchL, BchO, BchM, BchF from *Roseobacter denitrificans* OCh114). For genomes containing the full set, the fourteen proteins were concatenated, the sequences were aligned using *muscle* 3.8.31 [22] and a phylogenetic tree was constructed using *iqtree* 2.1.1 [23] with default parameters (evolution model chosen automatically). In parallel, the homologs of a set of conserved ribosomal proteins, elongation factors and gyrases (L2, L3, L4, L5, L6, L14, L15, L16, L17, L18, L22, L23, L24, L29, L30, L36, S3, S4, S5, S7, S8, S10, S11, S12, S13, S14, S17, S19, Tu, G, GyrA, and GyrB from *Pseudomonas aeruginosa* PAO1) were retrieved from the same proteomes and a phylogenetic tree was constructed following the same procedure. The Erythrobacteraceae ribosomal and superoperon trees were constructed in a similar way; however, since the genomes available on NCBI repository are not always complete, alignments had to be curated semi-automatically to avoid artefacts due to missing proteins. The 16S rRNA gene tree of the strains most closely related to *Porphyrobacter* sp. ULC335 was extracted from a tree comprising all Erythrobacteraceae plus *Caulobacter crescentus* NA1000 (outgroup); the 16S rRNA gen was extracted using *rnammer* 1.2 [24]; the nucleotide sequence alignment was done using *muscle* 3.8.31 [22] and a phylogenetic tree was constructed using *iqtree* 2.1.1 [23] with default parameters (evolution model chosen automatically) and 1000 boostrap. The DDH (digital DNA-DNA hybridization) was estimated using the digital Genome-to-Genome Distance Calculator algorithm recommended by the Leibniz Institute DSMZ (Formula 2) [7]. The ANI (average nucleotide identify) as calculated using PYANI [6].

SFigure 3. Strains were considered to encode the anoxygenic phototrophy superoperon if they gave positive *blastp* matches with at least 13 out of the 14 proteins listed above. The strains were assigned to a genus and class based on NCBI classification scheme.

#### Analysis of the ULC335 community

SFile 1. The trace sequences from the ULC335 metagenome (SRR6814902) were downloaded from NCBI repositories, trimmed using *fastp* [15] reassembled using *metaspades* with standard parameters [25], and resulting contigs were classified using the *CLASSIFIER* software (http://rdp.cme.msu.edu/classifier/classifier.jsp)[26].

#### Flow cytometry and fluorescence microscopy

##### Live-dead staining

Cells from 500µl of culture were resuspended in the same volume of NaCl 0.9% and incubated in the dark for 30 minutes after the addition of 1µl Invitrogen Baclite Live/Dead stain (mixture of equal volumes of components A and B). Cells were analyzed by flow cytometry and fluorescence microscopy. For the flow cytometry, cells were analyzed on a BD Accuri C6 Plus Flow Cytometer from a 1/100 dilution in NaCl 0.9%. Thresholds: 500 on FSC-H and 6000 on SSC-H. The zones corresponding to the dead, compromised, and live subpopulations were determined empirically based on the density distribution of live and 2.5% glutaraldehyde-fixed cells on a FL1 (green; excitation at 488nm, 533/30 emission filter) and FL3 (red; excitation at 488nm, 670LP emission filter) scatter plot. Visualization and quantification was done with R [27] using the *flowCore* [28] library after margin and doublet events removal using *PeacoQC* [29]. The same cells were analyzed using fluorescence microscopy (Leica DM4 B, Leica DFC7000 T camera, 100x objective : Leica 1.25 oil PH3 (506313)) with GFP and RFP settings; cells on phase-contrast, GFP, and RFP images were counted with FIJI [30] using a customized semi-automated script; analysis and plotting was performed with R base functions. Only dead and live subpopulations could be distinguished with fluorescence microscopy; a high-GFP and high-RFP signal “compromised” subpopulation could not be reliably identified, presumably because of the different optical settings and detection limits of flow cytometry and fluorescence microscopy.

##### DyeCycle Orange staining

Cells from 500µl of culture were resuspended in the same volume of NaCl 0.9%, fixed with 2.5% glutaraldehyde for 15 minutes, and incubated in the dark for 30 minutes after the addition of 1µl Invitrogen DyeCycle Orange stain. Cells were analyzed by flow cytometry on a BD Accuri C6 Plus from a 1/100 dilution in NaCl 0.9%. Thresholds: 500 FSC and 6000 SSC. Subpopulation analysis was done using R with base functions. The coordinates of the highest density in the regions of the 1N and 2N peaks were determined from the density of a 1024-break histogram of the FL1 fluorescence using the *density* function. The distance between the 1N and the 2N coordinates was used to predict the theoretical position of 3N and 4N peaks. The best combination of underlying 1N, 2N, 3N, and 4N logistic density distributions explaining the empirical density distribution was determined by tuning the scale factors. The lowest density point between the 1N and the 2N peaks on the empirical density plot was used as a proxy for the “actively replicating” population, the assumption being that high replication rates would lead to a continuum between the 1N and 2N peaks and the minimum between the two could be used as a replication index.

#### Survival and competition experiments

##### Bacteriochlorophyll *a* content depending on the growth phase

Strains of interest were grown in BG11-P medium for approximately 60h at 28°C with a 170 rpm shaking, in dark conditions. Cultures were adjusted to OD650 = 0.001 in BG11-P medium and distributed in a 24-well plate: 12 wells per strain containing 1.3 ml of culture. Three identical plates were prepared for the three light conditions tested: continuous light, continuous dark and 12h/12h dark-light cycles. Plates were incubated at 28°C with shaking. After 18h, 24h, 48h, 72h and 96h of growth, the cultures of two wells per strains were collected on each plate and OD650 measured in a SpectraMax i3x plate reader (Molecular Devices). 900µl of culture were centrifuged and pellets were resuspended in 1ml of a 0.9% NaCl solution. After centrifugation, pellets were resuspended in 250µl of methanol, centrifuged again and the clear supernatant was transferred in a new tube. Tubes were kept in the fridge and protected from light. 200µl of supernatant were loaded in a 96-well plate with flat bottom. Absorbance at 771 nm was measured to evaluate the production of bacteriochlorophyll a by the strains of interest in the different conditions. A full spectrum from 350 to 1000 nm (every 2 nm) was also recorded to evaluate the presence of other pigments. Measures were acquired in a SpectraMax i3x plate reader; samples were proceeded in two sets of 6 to avoid evaporation. The relative amounts of Bchl*a* and carotenoids (460 and 480 nm) were obtained by dividing the absorbances by the absorbance for the wild type DL condition (reference).

##### Survival depending on the genotype and the culture conditions

Late exponential phase cultures of the strains of interest (wild type, B^-^C^+^, B^++^C^++^) were diluted to OD650=0.001 in BG11-P and grown in 24-well plates (28°C, 170 rpm) under continuous light, continuous dark, or 12h/12h dark-light cycles in triplicates. At the beginning of stationary phase (at 48h), cultures were homogenized by pipetting and 100 µl of a 1.6*10^-5^ dilution of each replicate were plated on BG11-P; the colony forming units (CFUs) were calculated from the plate counts. After four additional days spent in stationary phase (at 144h), the procedure was repeated and CFUs were calculated again. The CFUs at 48h was used as a baseline and survival was calculated as the percent remaining cells at 144h.

##### Competition experiments

Late exponential phase cultures of the two mutant strains (B^-^C^+^ and B^++^C^++^) were mixed with the wild type strain in equal proportions (OD650=0.001) and grown in 24-well plates (28°C, 170 rpm) under continuous light, continuous dark, or 12h/12h dark-light cycles in triplicates. The dual cultures were plated at 48h and 144h like for the survival experiment, and the ratios of the mutant to the wild type were calculated for both timepoints based on CFUs.

#### Primers used in this study

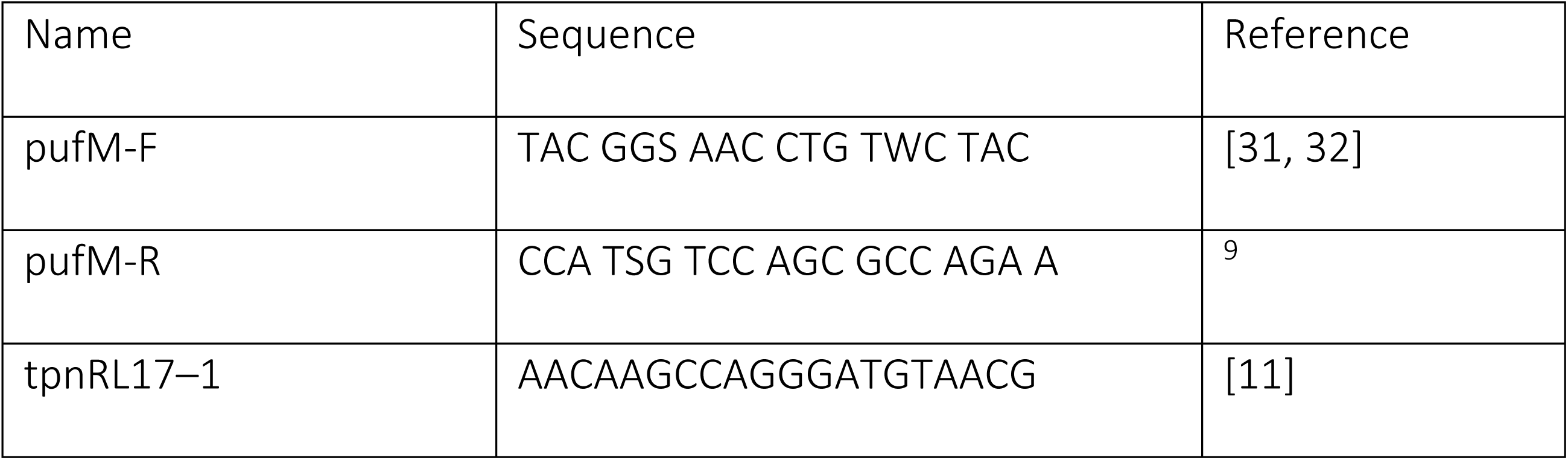

### QUANTIFICATION AND STATISTICAL ANALYSIS

Details on the quantification and statistical analyses performed can be found in the relevant subsections of the “METHOD DETAILS” and in the figure legends.

### ADDITIONAL RESOURCES

None.

